# Gene content of the fish-hunting cone snail *Conus consors*

**DOI:** 10.1101/590695

**Authors:** Reidar Andreson, Märt Roosaare, Lauris Kaplinski, Silja Laht, Triinu Kõressaar, Maarja Lepamets, Age Brauer, Viktorija Kukuškina, Maido Remm

## Abstract

**Background:** *Conus consors* is a fish-hunting cone snail that lives in the tropical waters of the Indo-Pacific region. Cone snails have attracted scientific interest for the amazing potency of their venom, which consists of a complex mixture of small proteins known as conopeptides, many of which act as ion channel and receptor modulators with high selectivity.

**Results:** We have analysed publicly available transcriptomic sequences from 8 tissues of *Conus consors* and complemented the transcriptome data with the data from genomic DNA reads. We identified 17,715 full-length protein sequences from the transcriptome. In addition, we predicted 168 full-length or partial conopeptide sequences and characterized gene structures of several conopeptide superfamilies.

## Introduction

*Conus consors* is a marine gastropod of the species-rich and highly diverse Mollusca phylum and we present the first extensive study of this organism from a genomic point of view. The first few genomes from this phylum (California sea hare, pearl oyster, Pacific oyster, owl limpet, octopus, and a freshwater snail) have only recently been sequenced (Takeuchi et al., 2012; Zhang et al., 2012; Simakov et al., 2013; Albertin et al., 2015; Adema et al., 2017) The phylogenetic position of *C. consors* is provided in Figure 1, which was constructed with particular reference to the other mollusc species for which genomic data are available.

*C. consors* is a member of the *Conoidea* superfamily that consists of more than 700 species worldwide (Puillandre et al., 2014; Lavergne et al., 2015; Gao et al., 2017). *C. consors* lives in the tropical waters of the Indo-Pacific, inhabits sub-tidal coastlines, but is also found at depths of up to 200 meters, where it buries itself under sand and silt for shelter (http://biology.burke.washington.edu/conus/).

**Figure 1.**
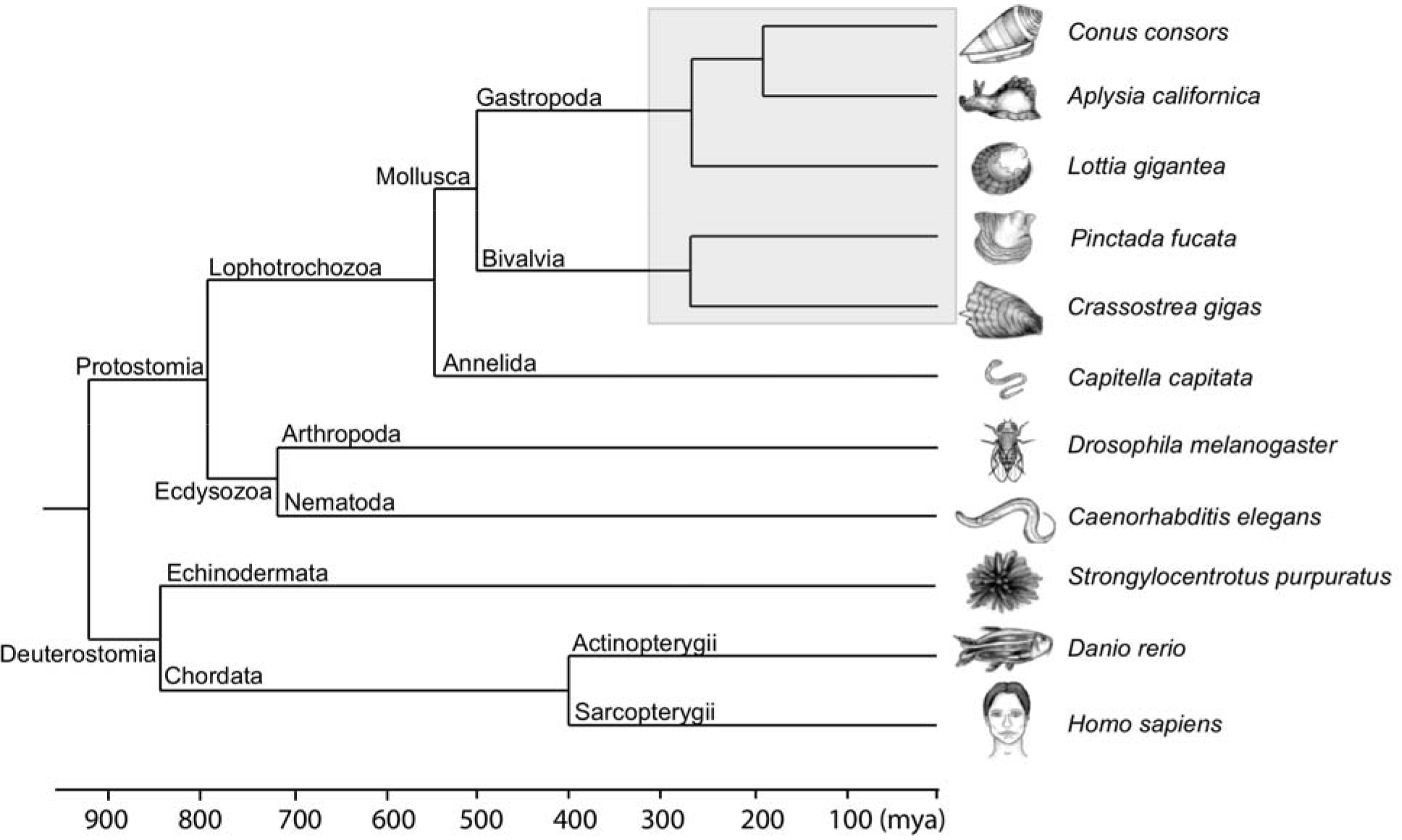
Phylogenetic position of *C. consors* in relation to some model organisms with sequenced genomes. The divergence times were obtained from the "Timetree of life" project (Hedges et al., 2015). Phylogenetic relationships within the *Mollusca* phylum are based on (Smith et al., 2011b) and (Kocot et al., 2011). The nodes included in the grey box are not time-scaled.

The cone snails have attracted scientific interest because of their pharmacologically active venom, which may provide leads in the search for novel drugs. The venom is a complex mixture of small peptides, termed conopeptides, that primarily act as ion channel modulators (Han et al., 2008; Favreau & Stöcklin, 2009; Lewis et al., 2012; Neves et al., 2015; Mir et al., 2016; Liu et al., 2018). When *C. consors* injects a fish with its venom, the fish is paralyzed within a few seconds and secured *via* a harpoon-like device. This “hook and line” strategy (Olivera, 1997) is unique to cone snails and makes up for their inability to chase prey.

Previous peptidomic and proteomic studies have revealed that the venom of cone snails is a complex mixture of several hundred peptides that shows both inter- and intra-species specific variability (Biass et al., 2009; Dutertre et al., 2010, 2013, 2014; Abdel-Rahman et al., 2011; Fu et al., 2018). Some variations in venom properties are linked to predation or defence stimuli (Dutertre et al., 2014).

To gain insight into the complexity of *C. consors*, we analysed transcriptome and genome sequences with the focus on gene content.

## Materials and Methods

### Transcriptome assembly

For assembly, we used publicly available sequencing reads generated by the CONCO consortium (Project #PRJNA271554 at NCBI SRA database). The transcriptome assembly included three steps: pre-processing of raw reads, separate assembly of tissue-specific transcriptomes from eight different tissues (venom duct, salivary gland, nerve ganglion, osphradium, mantle, foot, proboscis, and venom bulb) and combining transcriptomes into one non-redundant transcriptome set.

For pre-processing we trimmed the low quality 3’-ends of Illumina paired-end reads with the FASTQ Quality Trimmer from the FASTX Toolkit package version 0.0.13 (http://hannonlab.cshl.edu/fastx_toolkit/) using the quality cut-off (“-t”) at 30 and set the minimum length of the reads (“-l”) at 50 bp. We cleaned the reads with DeconSeq 0.4.1 (Schmieder & Edwards, 2011) and almost 850 million reads remained (in total ~800 Gbps).

For assembly of the transcriptome, we used the Trinity assembler (version 2012-06-08) (Grabherr et al., 2011) to create *de novo* transcripts for each sample with a minimum assembled contig length (“--min_contig_length”) set to 201 nucleotides.

Finally, in order to obtain a non-redundant set of sequences, we clustered the transcripts with CD-HIT-EST (Li & Godzik, 2006) using a sequence identity threshold (“-c”) of 0.98. The clustered transcriptome set is called the TRINITY transcriptome.

### Genome assembly

We have used publicly available sequencing reads generated by the CONCO consortium using a Roche 454 Genome Sequencer and an Illumina/Solexa GAII (Project #PRJNA267645 at NCBI SRA database). The average lengths of Roche 454 and Illumina reads were 354 bp and 104 bp, respectively. Four different types of data were used for the genome assembly: Roche 454 shotgun-sequenced reads, artificial 454 reads from an Illumina preliminary assembly with SOAPdenovo, six libraries of Illumina paired-end reads (300 bp and 600 bp insert sizes), and three libraries of Illumina mate pair reads (1.2 kbp, 3 kbp, and 7 kbp insert sizes). Detailed specifications for these libraries are provided in Supplemental Article S1.

During pre-processing, low quality 3’ ends of Roche 454 and Illumina reads were trimmed with the FASTQ Quality Trimmer. A quality cut-off (“-t”) was set to 30 and the minimum length of the reads (“-l”) was set to 50 bp. Consequently, reads were cleaned of human and bacterial contamination with DeconSeq 0.4.1. Identity (“-i”) and coverage (“-c”) cut-offs of 90% were used when scanning reads against human genome NCBI GRCh37 patch release 8 and 2,370 different bacterial strains. For the third step, SeqClean (version 2011-02-22) (https://sourceforge.net/projects/seqclean/) was used to remove any vector contaminations, linkers or adapter sequences. Tool was executed with default parameters excepting a minimum length of valid reads (”-l 50”), trimming of polyA/T tails, and low-complexity screening was disabled (”-A -L”). Reads were scanned against UniVec database build 7.0 (http://www.ncbi.nlm.nih.gov/tools/vecscreen/univec/) to remove any vector sequences.

Assembly included two distinct steps. At first, SOAPdenovo 2.04 (Luo et al., 2012) was used to create the initial genome assembly with Illumina paired-end/mate-pair reads. The goal was to create 454 “pseudo-reads” from the Illumina assembly as additional input data for Newbler. SOAPdenovo was applied with a k-mer word size of 37. The SOAPdenovo assembly generated many scaffolds that contained unresolved gaps (strings of “N”s). These scaffolds were split into 300 bp long sub-sequences with 200 bp overlaps to eliminate incorrect estimation of gap sizes using EMBOSS splitter (Rice, Longden & Bleasby, 2000). As a second step, Newbler 2.7 (https://sequencing.roche.com/) was run with the parameters “-large -rip -mi 98 -ml 100” to assemble all three types of reads – 454 (maximum read length 1,892 bp), “pseudo” 454 (300 bp) and Illumina (145 bp) – into one unique dataset. Contigs longer than 200 bp were reported in final assembly.

### Discovery of full-length genes from the transcriptome

We compiled a list of full-length genes from the TRINITY transcriptome using the following criteria:

1. We selected transcripts that exhibit at least 95% of their length matched to the genome using a BLASTN (version 2.2.22) (Altschul et al., 1997) alignment search. We performed unique mapping by first finding pairwise alignments between a transcript and a genomic region where the given alignment had the highest homology bitscore for both the transcript and genomic regions (seeds). For each seed we added the alignments for which the same transcript had highest alignment bitscore with the given genomic regions.
2. We annotated these transcripts using a BLASTX homology search against the UniRef100 database (Nov. 15, 2013) (Suzek et al., 2007). When homology to a given protein reached at least 75%, we annotated the transcript with its putative corresponding protein. In cases where there were multiple candidate proteins, we chose the one with highest cumulative alignment bitscore.
3. The cumulative bitscore of all transcript alignments with a given protein had to be greater than or equal to 100 bits.
4. All partial transcript homologies with a given protein had to be in the same translational frame.
5. The Open Reading Frame (ORF) had to be in one single translational frame, i.e. both the start and stop codons were present in the same frame.
6. The ORF start codon had to be located no more than 10 amino acids after the start of the first alignment and the stop codon not more than 10 amino acids before the end of the last alignment.

In cases where all of these criteria were met, we assigned the protein from the UniRef100 database as the annotation of a given transcript and generated the predicted protein sequence from the ORF.

### Annotation of conopeptides

We used four approaches to annotate conopeptide sequences from the assembled genome: 1) a BLAST search against the UniProtKB/Swiss-Prot database (release 2012_10) (The UniProt Consortium, 2015); 2) a HMM search using software HMMER 3.0 (http://hmmer.org/) (Eddy, 2011) against conopeptide HMM profiles (Laht et al., 2012); 3) a BLAST search against peptide sequences from *C. consors* venom proteomic data (Violette et al., 2012); and 4) a BLAST search against conopeptide sequences predicted from the transcriptome data of *C. consors*. In all four cases we applied an E-value cut-off of 10^−5^. We ran the HMMER and BLAST searches with default parameter values, except that we turned off the BLAST filtering option (-F F). We discarded matches that covered less than 50% of the length of their respective HMM profiles. We manually assessed the alignments and domain boundaries for all predictions.

### Data availability

Draft genome assembly of the cone snail can be retrieved from the GenBank database with following assembly ID: GCA_004193615. Gene and protein sequences predicted from transcriptome are included in Supplemental Data S4 (in FASTA format).

## Results and Discussion

### Transcriptome and genome assembly

Transcriptome assemblies were created with Trinity software using read libraries from eight different tissues (venom duct, salivary gland, nerve ganglion, osphradium, mantle, foot, proboscis, and venom bulb). The total number of transcripts (including isoforms) was 1,535,709 and ranged from 85,807 (“Foot” sample) to 240,307 (“Mantle” sample) and contained around 1,062 Gbp of sequence. The average length of the resulting transcripts for all samples was 692 bp, N50 = 2,452 bp, and the longest sequence was 29,867 bp. After clustering the results from eight samples with CD-HIT-EST, the final dataset contains 587,852 transcripts (~324 Gbp in total). The transcriptome data was used to compile a full-length gene list and to predict conopeptide genes.

For genome assembly we used a strategy similar to the one employed to assemble the genome of the fire ant *Solenopsis invicta* (Wurm et al., 2011). Briefly, this strategy consisted of two major steps: (a) assembly of Illumina reads (9 libraries, overall 51 Gbp of raw data) into larger contigs using SOAPdenovo software and (b) combining the resulting Illumina contigs and original paired-end reads from the Illumina and unpaired reads from Roche 454 libraries (1 fragment library, overall 6 Gbp of raw data) into a final assembly using the software Newbler (Supplemental Article S1 Figure 1). The assembly of Illumina reads into longer artificial reads was required because Newbler is not optimized to work with short Illumina reads. In step (b), the original Illumina reads were also included to provide additional information about the distance between paired reads.

The final assembly of *Conus consors* genomic reads resulted in a 2,049 Mbp sequence consisting of 2,688,687 scaffolds and contigs with an N50 size of 1,128 bp. Newbler software is able to estimate the size of the entire genome based on k-mer frequency distribution. *C. consors* genome was estimated to be 3.025 Gbp, which is within the range of other cone snail genomes (http://genomesize.com/). The genomic DNA resulting from this assembly is fragmentary; nevertheless, the protein-coding exons are generally contiguous. Therefore, we were able to use it as an additional source of information in gene prediction process and for characterization of conopeptide gene structures.

The genome of *C. consors* is rich in repeats. Approximately 49% of the genome sequence contains repeated sequences, half of which are low-complexity (mononucleotide, dinucleotide, trinucleotide and tetranucleotide) repeat elements. Detailed analysis of repeat elements present in the genome is shown in Supplemental Article S1.

### Coverage of core genes in transcriptome and genome

To evaluate the completeness of our transcriptome and genome assemblies we calculated the length coverage of core genes from the Core Eukaryotic Genes Mapping Approach (CEGMA) dataset (Parra, Bradnam & Korf, 2007; Parra et al., 2009). This dataset consists of 458 core proteins that are universally present in 6 eukaryotic species: *Homo sapiens, Drosophila melanogaster, Arabidopsis thaliana, Caenorhabditis elegans, Saccharomyces cerevisiae* and *Schizosaccharomyces pombe.* A similar method has previously been used to evaluate the quality of two different ant genome assemblies (Smith *et al.* 2011; Wurm *et al.* 2011). Coverage (fraction of amino acids detected by TBLASTN search using core protein dataset as a query) of core genes in our transcriptome and genome data is shown in Figure 2. The median coverage of core genes is 99.7% for transcriptome and 93.4% for the genome. Similar genome coverage was observed for other mollusc genomes (Supplemental Article S1). One has to take into account that TBLASTN is somewhat limited in finding short exons in genome, thus the coverage of core genes measured from genome will always be lower than coverage in transcriptome. An illustration of core gene alignment from *C. consors* genome is shown in Figure 3.

**Figure 2.**
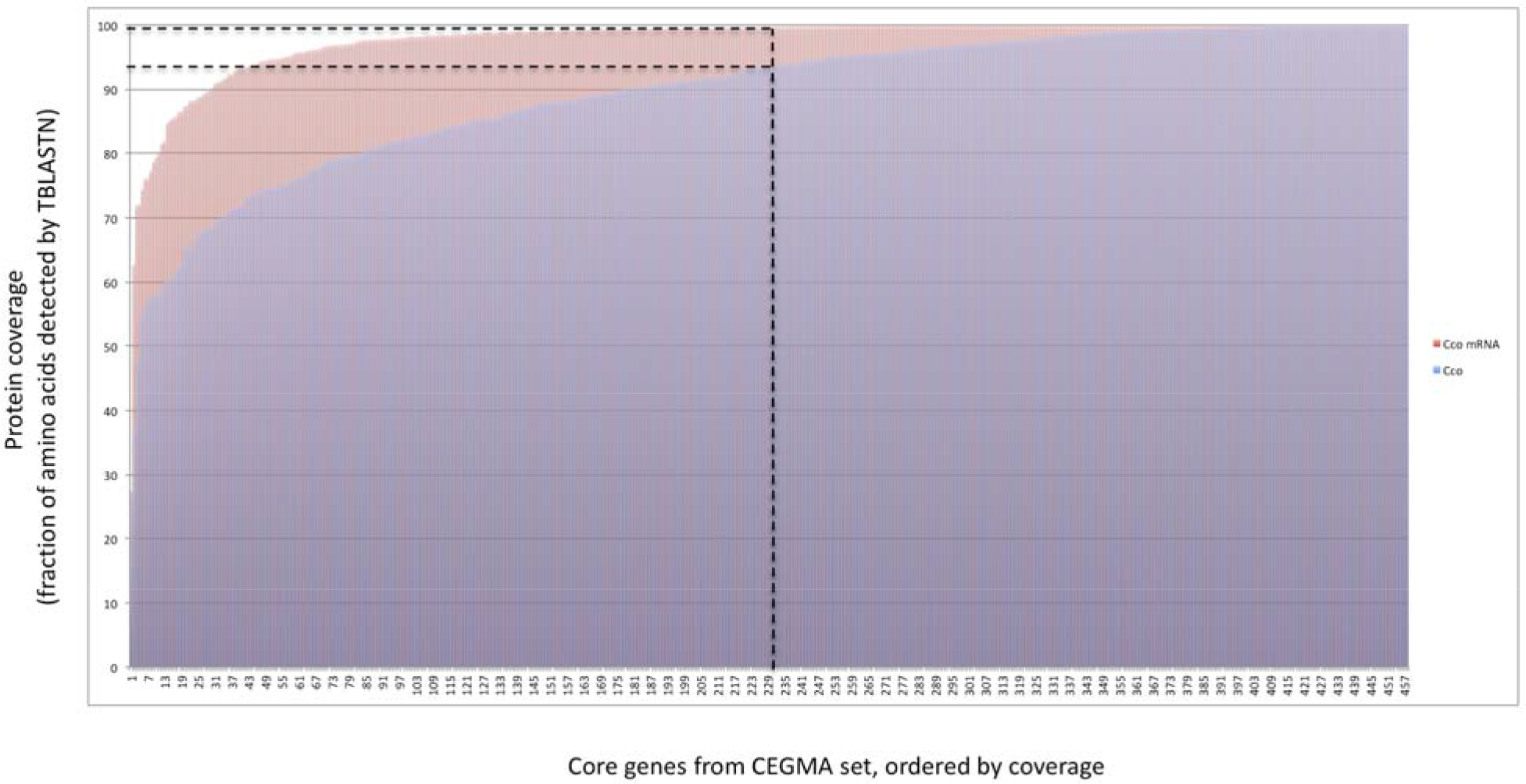
Coverage of 458 core proteins from the CEGMA dataset in *C. consors* transcriptome and genome. Coverage is defined as fraction of amino acids detected by TBLASTN search using core protein dataset as a query.

### Gene content of *C. consors*

We predicted full-length protein sequences from the transcriptome data using a reciprocal homology search between the transcriptome and the UniRef100 protein database. The genome sequence was used to confirm the existence of genes predicted from transcriptome. We consider the resulting 17,715 full-length proteins to be a reliable prediction of protein-coding sequences of *C. consors*. The collection of mRNAs and translated protein sequences in FASTA format is available in Supplemental Data S4. It has to be kept in mind that the actual number of protein-coding genes is somewhat larger due to the fact that transcriptome analysis cannot reveal genes that are expressed at low levels, in other tissues or just temporarily.

**Figure 3.**
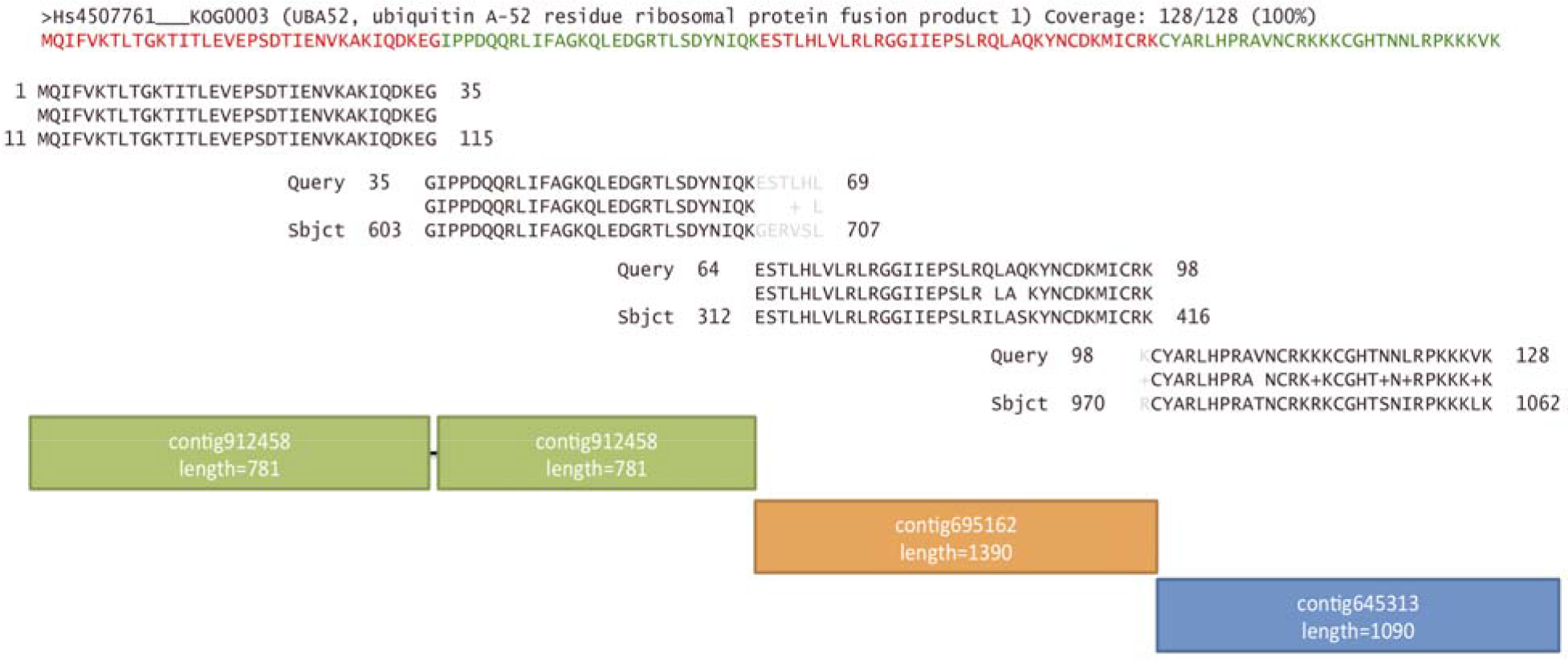
Example of gene content in the genome. TBLASTN against the genome using CEGMA (core protein set present in all eukaryotes) protein Hs4507761 as a query. Red and green text denote location of alternating exons in the human gene *UBA52*. Red orange and blue boxes are matching regions from contigs or scaffolds of the *C. consors* genome. Alignment between Hs4507761 and translated genomic DNA is shown in the middle.

We analysed the number of tRNA, rRNA and of other non-protein-coding genes using tRNAScan-SE (Lowe & Eddy, 1997) and Infernal software (Nawrocki, Kolbe & Eddy, 2009). We detected a total of 761 different tRNA genes in the *C. consors* genome, 2500 miRNA genes and many other types of RNA genes. Detailed analysis of RNA genes present in the genome is shown in Supplemental Article S1 and full list of detected RNA genes is shown in Supplemental Article S1 Table 2.

### Conopeptide sequences

To identify conopeptide sequences in the transcriptome and genome of *C. consors*, we used several sources of data with previously known conopeptide sequences or hidden Markov model (HMM) profiles. Conopeptide sequences available in the UniProtKB/Swiss-Prot database (975 peptides from more than 30 different superfamilies), 64 conopeptide hidden Markov model (HMM) profiles from 20 different superfamilies (Laht et al., 2012), 126 peptide sequences from the *C. consors* proteome sequencing (Violette et al., 2012), and conopeptide precursor sequences predicted from the transcriptome data (135 distinct precursor sequences from 23 different superfamilies) were used. In addition to main transcriptome data we also used another dataset (CC8 transcriptome), sequenced earlier. This additional transcriptome data originated from two ESTs libraries constructed from venom duct and salivary gland tissues. The procedure for obtaining CC8 transcriptome sequences is described in (Terrat et al., 2012). The genome sequence was also checked for potential conopeptide genes in hope that it complements transcriptome-based data.

To estimate the overall number of conopeptides encoded by *C. consors*, we aligned predicted protein sequences obtained from the genome, transcriptome, and proteome into multiple alignments (Supplemental Data S3.). Sequences from different datasets exhibit clear clusters with slight variations between individual sequences. Closely related sequences were merged into clusters if the difference between sequences did not exceed 4 amino acids and the overall number of sequence clusters was counted. Example of multiple alignment of sequences from the O1-superfamily is shown in Figure 4. This way we estimated that *C. consors* could have at least 168 conopeptides: 27 with previously known sequence and 141 novel sequences. In addition, we list 46 dubious sequences, which were only detected in the genome and did not have any closely related sequence in databases. These might be products of pseudogenes, products of wrongly predicted genes or peptides with other functions. However, it is not excluded that some of these “dubious” clusters might represent novel conopeptides. The superfamilies M, O1, and A comprise about 42% of all identified conopeptides in the *C. consors* (Table 1), which is in concordance with previously published data (Puillandre et al., 2012).

**Figure 4.**
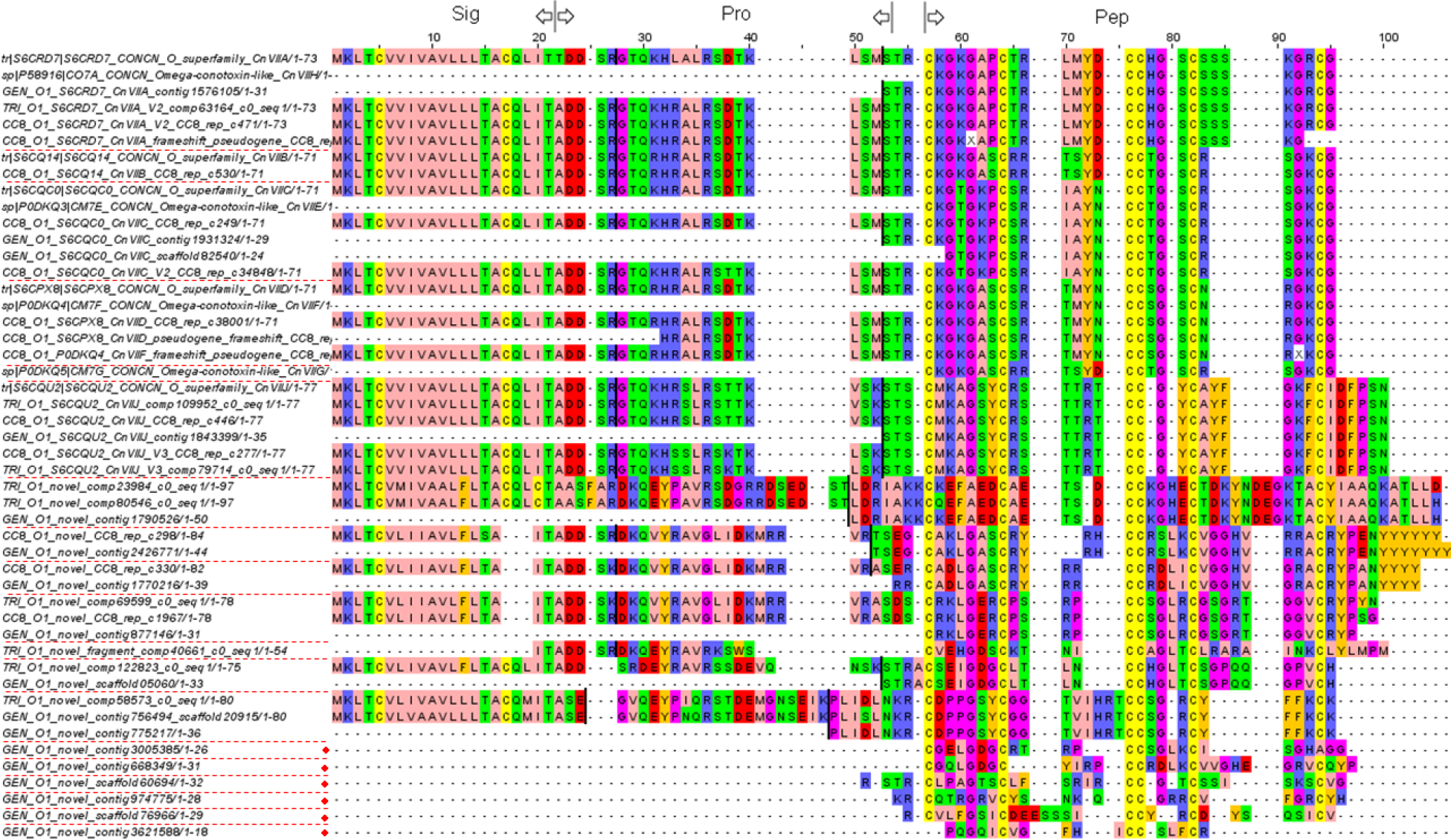
Example of conopeptide gene clusters. A subset of O1 superfamily gene clusters is shown. Red lines denote boundaries of gene clusters. Red dots indicate “dubious” genes, which show some similarity with conopeptides, but are not counted as conopeptide genes.

**Table 1.**
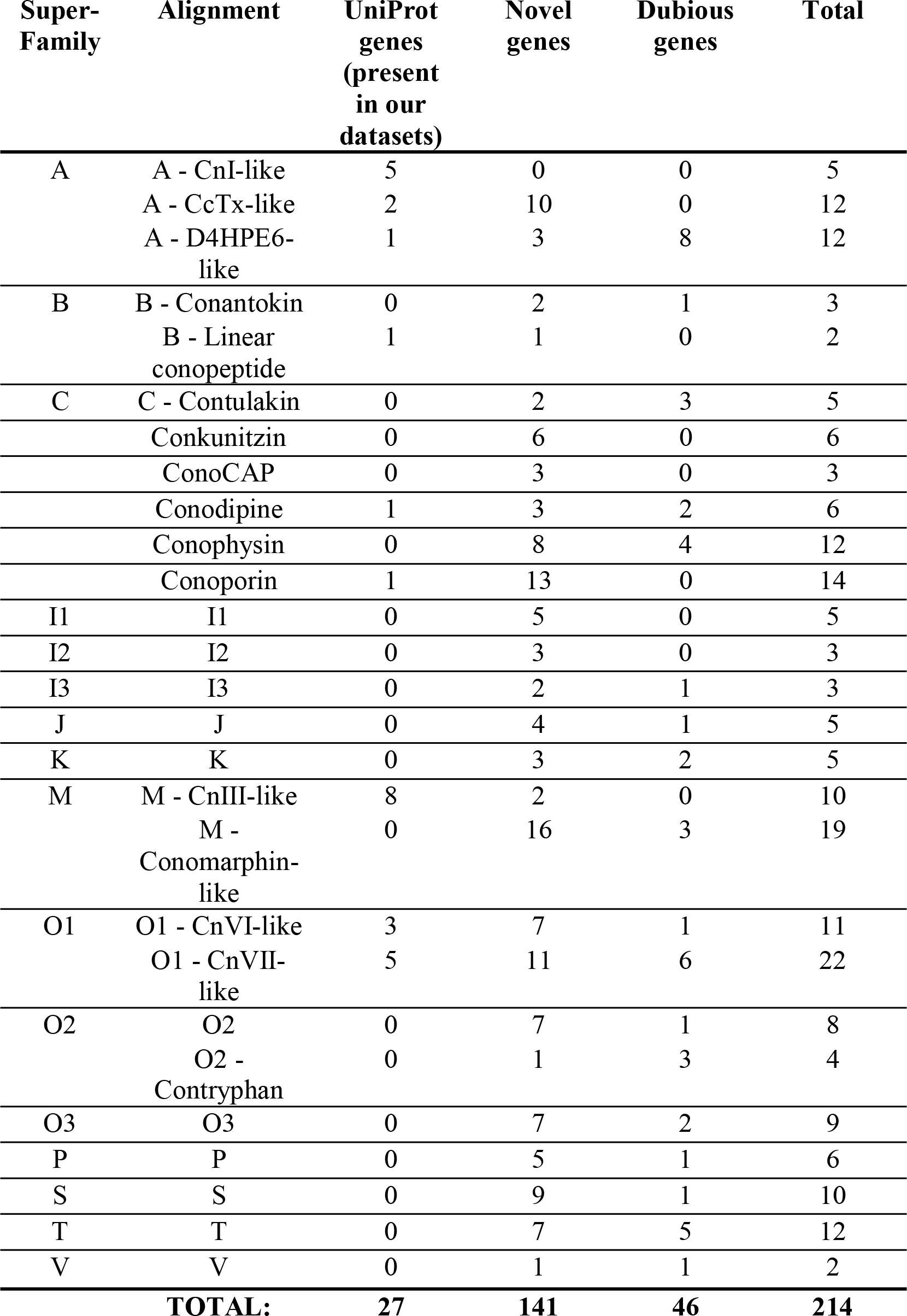
Number of conopeptide genes predicted from the *C. consors* genome and transcriptomes, ordered by superfamilies.

### Conopeptide genes in genome

The majority of conopeptide superfamilies are known to contain introns that separate different functional domains (Olivera et al., 1999). The genome sequence allows us to identify the genomic structure of some conopeptide genes. Sequences that code for signal, propeptide, and mature peptide domains were retrieved for 15 conopeptides from 14 superfamilies (Figure 5). It is noteworthy that we can identify several different exon-intron organizations within the conopeptide genes. The first exon of the most abundant type encodes for the complete signal peptide sequence together with a variable length fragment of a pro-peptide, while the first exon of genes encoding type A, I1, I3, and M conopeptides encode the entire signal sequence. Pro-peptides appear to be encoded by one, two, or three different exons. Only conodipine genes are devoid of pro-peptide sequences. Finally, in the unique case of J-conopeptides, their genes appear to be made of a unique encoding exon containing, successively, a signal, an N-terminus pro- and a mature peptide, followed by a C-terminus pro-sequence.

**Figure 5.**
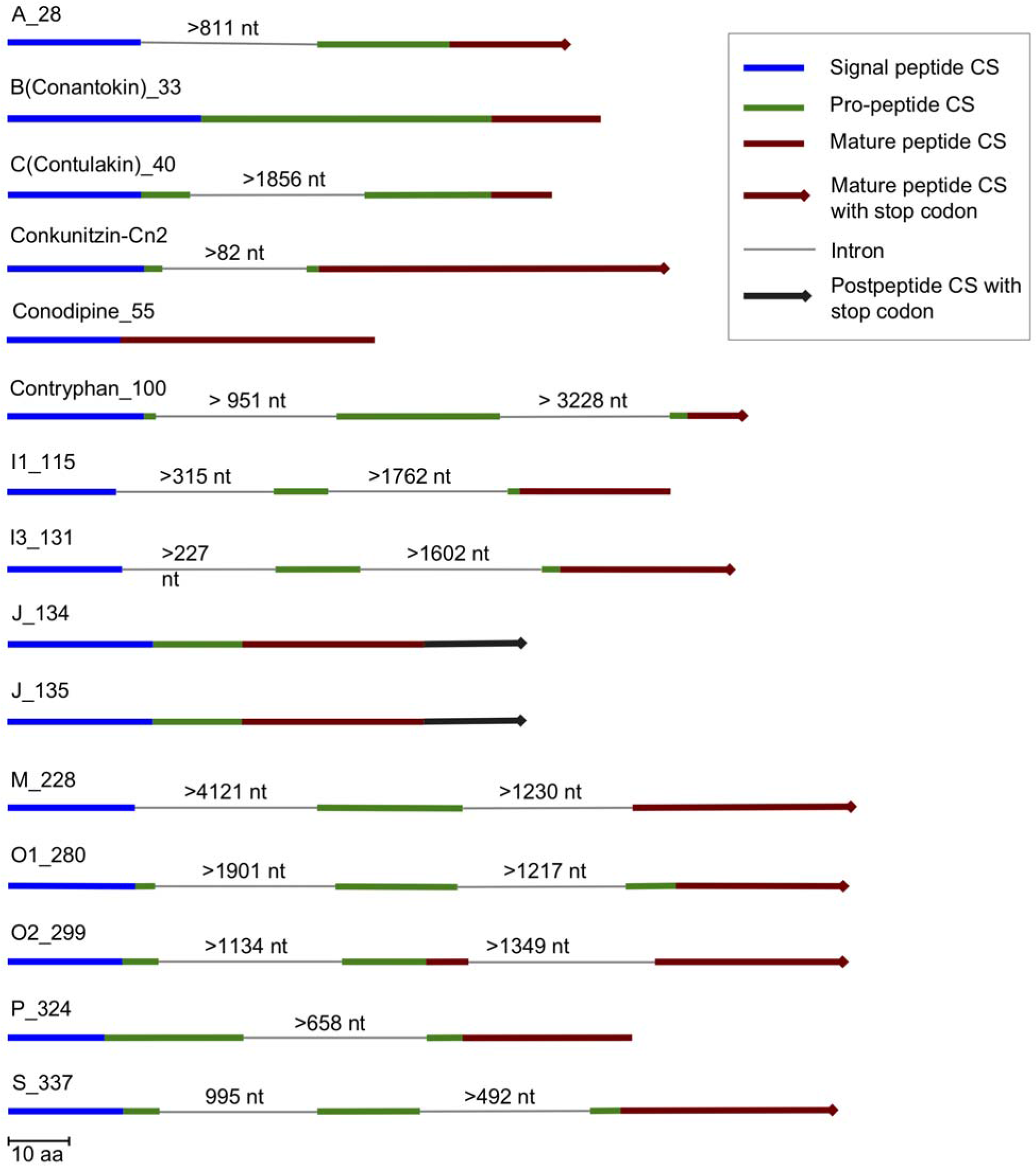
Conopeptide gene structures within the genome of *C. consors*. Each sample represents one conopeptide gene. The peptide coding sequences (CS) for signal, pro- and mature peptides are represented by bold blue, green, and red lines. The length of each line is proportional to the number of amino acids. The introns are represented as thin grey lines and the length of the intron sequences is indicated in nucleotides above each line. The symbol '>' indicates that this gene was not assembled into a single contig and that the intron length is therefore not precisely known. Sequences of the conopeptide genes and additional information are available in Supplementary Data S3.

## Conclusions

The annotation of a fish-hunting cone snail *C. consors* genome and transcriptome gives us a closer opportunity to peek into the complexity of its genes. The analysis of the combined eight different transcriptomic and genomic datasets resulted 17,715 full-length protein sequences. In addition, 168 conopeptide sequences were identified and in several cases the gene structures of conopeptide superfamilies were characterized. We have found several gene coding clusters that might represent novel conopeptides and are therefore good candidates for future studies.

## Supplemental information

The following additional data are available with the online version of this paper. Supplemental Article S1 contains a detailed description of all supplementary analysis and methods. Supplemental Data S2 contains list of predicted RNA genes, clustered by RFam category. Supplemental Data S3 contains alignments of conopeptides from each superfamily. Gene and protein sequences predicted from transcriptome are included in Supplemental Data S4 as two separate FASTA format files.

## Authors’ contributions

MRe and RA were responsible for drafting the manuscript. MRo, LK, SL, TK, AB, VK, RA and MRe analyzed the sequencing data. All authors have read and approved the final manuscript.

## Supporting information

Supplemental Article S1

## Acknowledgements

Tõnu Margus and Aleksander Sudakov gave advice on construction of phylogenetic trees and gene content analysis. We would like to thank Tim Stockwell, Philippe Favreau, Daniel Biass, Yves Terrat and Dusan Kordis for valuable discussions during the initial survey of the genome sequence data.

## Funding Statement

AB, SL, RA, LK, MRo, ML, TK, VK and MRe were supported by the EU FP6 CONCO project, SF0180026s09 and IUT34-11 from the Estonian Ministry of Education and Research and by the EU ERDF grant No. 2014-2020.4.01.15-0012 (Estonian Center of Excellence in Genomics and Translational Medicine). RA was also supported by the EU ERDF grant No. 2014-2020.4.01.16-0125. The computational analysis was partly carried out on the High Performance Computing Center of University of Tartu.

