## Supplemental Article S1 for "Gene content of the fish-hunting cone snail *Conus consors*"

### 1. GENOME

#### Raw data and assembly procedure

Data used for the genome assembly is shown in Supplementary Table 1. The assembly was performed in two distinct steps (Supplementary Figure 1). First, Illumina reads were assembled with SOAPdenovo 2.04 (Luo *et al.* 2012) to create 454 "pseudo-reads" as an additional input data for Newbler. This resulted in 6,850,185 sequences with a total length of 3,771,619,740 bp. Many of the longer scaffolds contained unresolved gaps (strings of "N"s) and were split into 300 bp long sub-sequences with 200 bp overlaps to eliminate incorrect estimation of gap sizes with the EMBOSS splitter (Rice *et al.* 2000). That resulted in 6,544,872 sequences (1,805,641,865 bp) in FASTA format. Second, all three types of reads – the 454 reads (maximum read length 1,892 bp), "pseudo" 454 reads generated in the first step (300 bp), and Illumina paired-end and mate pair reads (145 bp) were assembled into one dataset with Newbler 2.7 (https://sequencing.roche.com/). The final assembly contained 2,688,687 scaffolds and contigs and covered a total of 2.05 Gbp of genomic sequence.

**Supplementary table 1. Description of the data used for genome assembly**

| Library | Type | Insert size | Max read length | # of reads/pairs | # of bases |
|---|---|---|---|---|---|
| **454** | | | | | |
| Lib_1 | Single | NA | 1,892 | 19 M | 6 Gbp |
| **Illumina** | | | | | |
| Lib_2-5 | Paired-end | 300 | 145 | 73 M | 14 Gbp |
| Lib_6-7 | Paired-end | 600 | 145 | 25 M | 6 Gbp |
| Lib_8 | Mate pair | 1,200 | 100 | 20 M | 4 Gbp |
| Lib_9 | Mate pair | 3,000 | 100 | 37 M | 7 Gbp |
| Lib_10 | Mate pair | 7,000 | 100 | 111 M | 20 Gbp |
| | | | **Total overall:** | **285 M** | **57Gbp** |

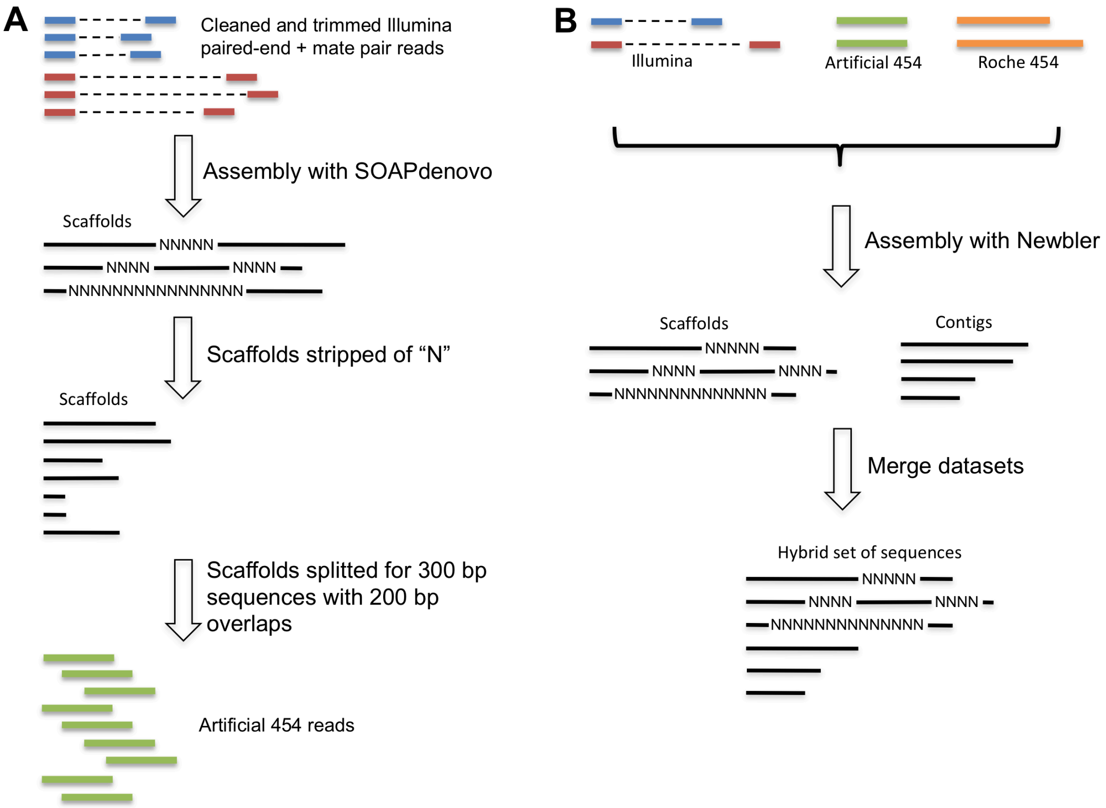


**Supplementary Figure 1. Genome assembly workflow.** First, a set of long overlapping reads was created using SOAPdenovo (A) and then all datasets were assembled with Newbler (B).

### Comparison with other mollusc genomes

Comparison of *C. consors* genome assembly with other mollusc genomes is shown in Supplementary Table 2. Although our assembly has rather low N50 value, we believe that protein-coding exons are well detectable in our data (see Supplementary Figure 2).

**Supplementary Table 2. Comparison of sequenced mollusc genomes. Sequencing and core gene coverage statistics of five mollusc species.** Raw sequence coverage of *C. consors*, *A. californica*, and *L. gigantea* was calculated by dividing the number of nucleotides in the raw reads with the estimated length of the genome. Coverage of core genes is expressed as fraction of amino acids detected from the genome by TBLASTN (Altschul *et al.* 1997) search.

| Species | Length of the genome (Mbp) | Raw sequence coverage | Length of assembled sequences (Mbp) | N50 (bp) | Median coverage of core genes (%) | Sequencing technology | Citation |
|---|---|---|---|---|---|---|---|
| *Conus consors* | 3,000 | 19x | 2,049 | 1,128 | 93.4 | Roche 454, Illumina | Current study |
| *Aplysia californica* | 1,800 | 10x | 716 | 264,327 | 91.6 | Sanger | Jessica Afoldi, personal communication |
| *Lottia gigantea* | 500 | 9x | 360 | 1,870,055 | 93.9 | Sanger | [1] |
| *Pinctada fucata* | 1,150 | 40x | 1,413 | 1,629 | 91.9 | Roche 454, Illumina | [2] |
| *Crassostrea gigas* | 600 | 690x | 559 | 19,400 | 93.5 | Illumina | [3] |

[1] Lottia genome data at JGI [http://genome.jgi-psf.org/Lotgi1/Lotgi1.download.ftp.html]
[2] Pinctada fucata genome ver 1.00 [http://marinegenomics.oist.jp/pearl/viewer/info?project_id=20]
[3] Zhang *et al.* 2012]

### Coverage of core genes in mollusc genomes

In order to test whether the fragmented genome assembly compromises gene discovery from the genome we compared *C. consors* core gene coverage (Parra *et al.* 2007, 2009) with coverage in other sequenced genomes. The results, shown in Supplementary Figure 2 indicate that core genes can be detected very similarly in different genomes. For example, *L. gigantea* sequence used in this analysis had N50 1,870,000 bp, due to the fact that it was sequenced with Sanger technology. This shows that coverage of protein coding-genes in mollusc genomes is not very sensitive to the N50 values and most protein-coding exons should be well represented in our assembly. It is likely that some genes in core set get lower coverage because they contain some rather short exons which are difficult to detect by TBLASTN analysis.

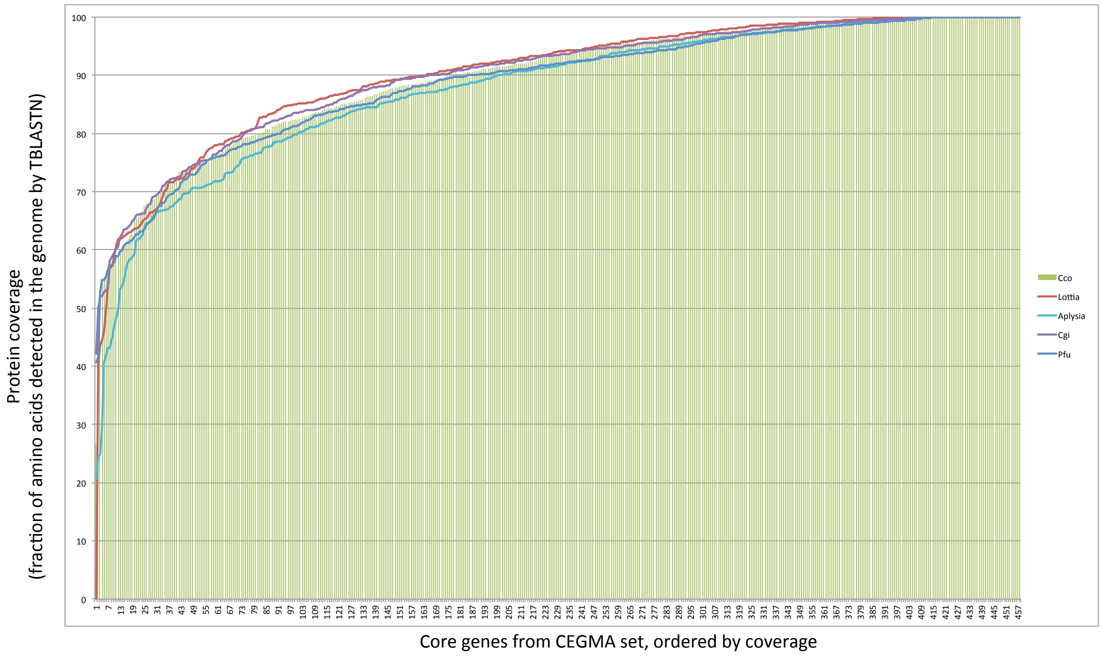


**Supplementary Figure 2. Coverage of 458 core genes from CEGMA core gene set.** Genes are ordered by their coverage in genomes. Colors indicate data from different species (green bars - *C. consors*, dark blue line - *P. fuctata*, red line - *L. gigantea*, light blue line - *A. californica*, magenta line - *C. gigas*).

### 2. REPEATS

#### Identification and classification of repeated regions in the genome

We used DUST (Morgulis *et al.* 2006), Tandem Repeats Finder (Benson 1999) RepeatMasker (http://www.repeatmasker.org) and RepeatScout (Price *et al.* 2005) software, in that order, to detect and classify repeated regions from the *C. consors* genome. Analysis was based on randomly selected 454 reads (5.7 million reads) instead of an assembled genome sequence, because assemblers often erroneously combine reads with similar sequences into one contig or fail to assemble repetitive regions at all. The proportions of different repeat types are shown in Supplementary Figure 3.

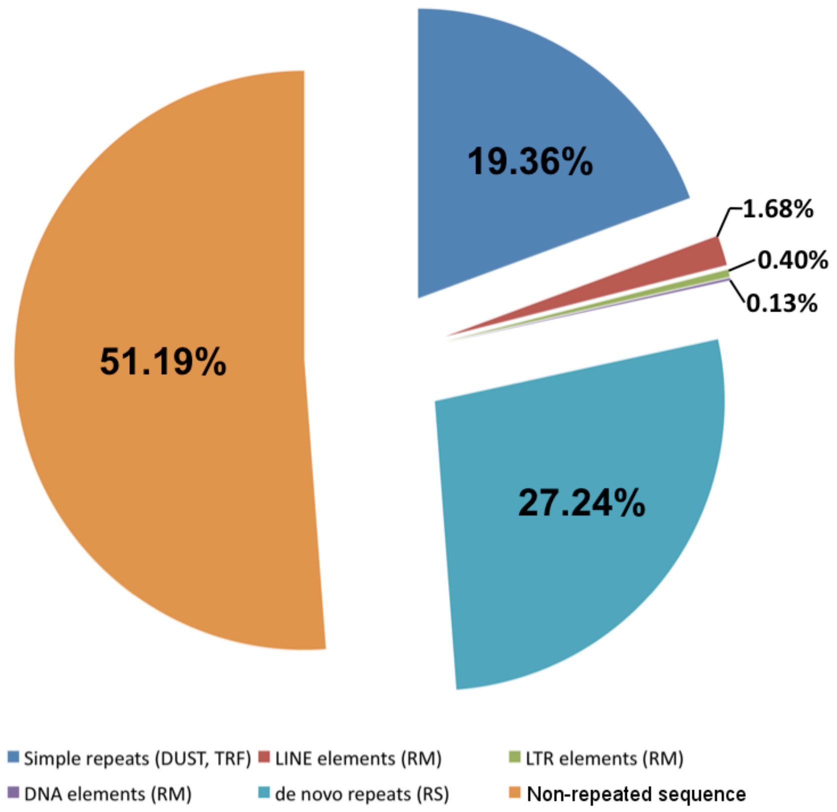


**Supplementary Figure 3. Proportions of the various repetitive element types and of the non-repeated sequences within the *C. consors* genome.** The repeated regions were detected by DUST, Tandem Repeats Finder (TRF), RepeatMasker (RM), and RepeatScout (RS), in that order. To avoid bias introduced by the assembly software, the fractions shown here were calculated from randomly selected 454 reads (in total >2 Gbp raw sequence).

Approximately 20% of the genome contains low-complexity (mononucleotide, dinucleotide, trinucleotide and tetranucleotide) repeats masked by both DUST and the Tandem Repeats Finder. About 2% of the genome is covered by known repeats that were identified by RepeatMasker and RepBase However, over 27% of the genome contains novel motifs that are not present in the current RepBase database and may represent potential mollusc-specific transposable elements such as DNA transposons and retrotransposons. In total, approximately 49% of the genome consists of repeats, which is comparable with the fraction of repeats within the human genome.

We have used a following procedure for masking repeats. At first, we detected low-complexity and tandem elements with DUST (cut-off score 15). Next, we used Tandem Repeats Finder 4.04 to locate tandem repeats with the following options: -r (no redundancy elimination), matching weight=2, mismatching penalty=2, indel penalty=7, match probability=80, indel probability=10, minimum alignment score to report=50, maximum period size to report=500. We then masked the previously described repeats with RepeatMasker 3.3.0, which includes RepBase 16.10 (Jurka *et al.* 2005) using these parameters: "-s -nolow –cutoff 400". Finally, we performed *de novo* repeat identification with RepeatScout 1.0.5. We used the assembled contigs to build an initial l-mer table using build_lmer_table ("-l 17") and then created a *de novo* repeat library with RepeatScout ("-l 17"). We masked the 454 reads with RepeatMasker ("-s -nolow -no_is -norna"), again using RepeatScout output as an input library.

### Comparison of repeat content with other mollusc genomes

The presence of *Conus consors* unknown repeat motifs in other mollusc genomes was studied to evaluate whether these are specific to *C. consors*. The fraction of repeats in other mollusc genomes is clearly lower - between 10% and 36% (Takeuchi *et al.* 2012; Zhang *et al.* 2012; Simakov *et al.* 2013). Additional analysis indicates that homologs of unknown repeats are mostly absent in other molluscs (Supplementary table 3). Thus, the higher fraction of repeats in the *C. consors* genome could be due to the appearance of new species-specific repeat families; however, a comprehensive analysis of these repeats is beyond the scope of the current study.

For this analysis, the RepeatScout repeat library was used with 11,155 unknown repeat types created for *C. consors*. Homologous regions of repeats were searched with BLAST (BLASTN with "–evalue 1e-03 –dust no" options) from four mollusc genome sequences using each mollusc genome as a query and recording only the best hits. Hits with less than 80% coverage of the *C. consors* repeat were discarded.

**Supplementary table 3. Presence of the unknown repeats within *Conus consors* in other mollusc genomes.**

| **Species** | **Number of *C. consors* motifs detected (total 11,155)** |
|---|---|
| *A. californica* | 305 (2.7%) |
| *P. fucata* | 171 (1.5%) |
| *C. gigas* | 143 (1.3%) |
| *L. gigantea* | 65 (0.6%) |

**Repeats in exon flanking regions**

We were able to confirm the higher frequency of repeated sequences in flanking regions of conopeptide gene exons. We tested whether low-complexity sequences (detected by DUST) are equally frequent in 50 bp neighbourhoods between conopeptide exons and all other gene exons. We detected low-complexity regions in the flanking regions of 14.4% (51/355) conopeptide exons and in the flanking regions of 9.2% (73,010/789,402) exons of other genes. We also observed a similar frequency of low-complexity repeats (8.8%) in randomly selected 50 bp long genomic sequences. Thus, the conopeptide exons contain low-complexity repeats in their flanking regions 1.6x more frequently than exons from other genes. This difference is statistically significant in the Fisher exact test (P=0.0017). The relative richness of low-complexity repeats could be one of the mechanisms that drive the fast duplication of conopeptide exons or entire genes.

The following procedure was used to detect repeats in exon flanking regions. First, we used BLAST to search the genome for potential exons, which we defined as more than 50 nucleotides long DNA sequences that have at least 97% identical local alignment with one or more transcripts. We considered conopeptide transcripts and other transcripts as separate sets. Second, we located and extracted 50bp long flanking regions, when possible, from both sides of the exons. In cases where overlap occurred among potential exons, we only considered the flanking regions of the longest exon. Altogether, we collected 355 flanking regions for conopeptide exons and 798,402 flanking regions for other exons. We generated a background set of 300,000 50-nucleotide-long random genome sequences, which did not overlap with either the exons or their flanking regions. We detected the low-complexity repeats in all sequence sets using the DUST software with a cut-off score of 30.

### 3. RNA GENES

#### Non-coding RNAs

We analysed the number of tRNA, rRNA and of other non-protein-coding genes using tRNAScan-SE (Lowe and Eddy 1997) and Infernal software (Nawrocki *et al.* 2009). We detected a total of 761 different tRNA genes in the *C. consors* genome, which is more than the estimated number in the human genome (513 different tRNA genes according to Genomic tRNA Database) (Chan and Lowe 2009).

Surprisingly, we also detected a large number of tRNA pseudogenes (21,972), which are classified as putative tRNA-derived short interspersed elements (SINEs) by the tRNAScan-SE software. tRNA-derived SINEs have been previously described in a number of eukaryotes (Ogiwara *et al.* 1999; Nishihara *et al.* 2006). For comparison, we searched both the human and mouse genomes using the tRNAScan-SE software with the same parameters and found 150 and 26,937 tRNA pseudogenes, respectively. Both of these values agree with previously published numbers of tRNA pseudogenes (Goodenbour and Pan 2006). The number of tRNA pseudogenes was also estimated in *Aplysia californica* and other mollusc genomes (Supplementary table 4).

**Supplementary table 4**. **Number of different tRNA gene sequences in compared genomes.**

| Species | Number of tRNA gene copies | Number of tRNA pseudogenes (putative tRNA-derived SINEs) |
|---|---|---|
| *Conus consors* | 761 | 21,972 |
| *Aplysia californica* | 274 | 17,347 |
| *Lottia gigantea* | 808 | 104 |
| *Pinctada fucata* | 903 | 4,345 |
| *Crassostrea gigas* | 431 | 1,974 |
| *Mus musculus* | 480 | 26,937 |
| *Homo sapiens* | 449 | 150 |

Infernal software was also used to detect other non-coding RNAs in the genome of *C. consors*. This analysis provided estimates of 2,546, 107, and 130 for the number of microRNA genes, snoRNA genes, and spliceosomal RNA genes, respectively (Supplementary file 2). MicroRNA is the most abundant type of non-coding RNA predicted with the most prolific microRNA gene being mir-598, which contains 1,516 sequences (Supplementary table 5).

**Supplementary table 5. MicroRNA genes predicted in *Conus consors* genome**. Rfam families with at least 10 predicted sequences are shown.

| Nr | Accession | Description | Count |
|---|---|---|---|
| 1 | RF01059 | microRNA mir-598 | 1,516 |
| 2 | RF00929 | microRNA mir-574 | 209 |
| 3 | RF01043 | microRNA MIR1023 | 89 |
| 4 | RF00885 | microRNA MIR821 | 67 |
| 5 | RF01005 | microRNA MIR530 | 51 |
| 6 | RF00673 | microRNA mir-217 | 42 |
| 7 | RF02244 | microRNA mir-785 | 32 |
| 8 | RF00920 | microRNA MIR444 | 30 |
| 9 | RF00452 | mir-172 microRNA precursor family | 29 |
| 10 | RF00690 | microRNA MIR408 | 28 |
| 11 | RF01021 | microRNA mir-558 | 28 |
| 12 | RF01911 | microRNA MIR2118 | 21 |
| 13 | RF01064 | microRNA mir-253 | 20 |
| 14 | RF00908 | microRNA MIR529 | 15 |
| 15 | RF00047 | mir-2 microRNA precursor | 15 |
| 16 | RF00756 | microRNA mir-299 | 14 |
| 17 | RF00884 | microRNA MIR815 | 14 |
| 18 | RF00704 | microRNA MIR397 | 13 |
| 19 | RF01943 | microRNA mir-999 | 12 |
| 20 | RF00660 | microRNA mir-214 | 11 |
| 21 | RF01061 | microRNA mir-548 | 11 |
| 22 | RF00714 | microRNA MIR535 | 10 |
