## Supplementary figures and images for "Gene content of the fish-hunting cone snail *Conus consors*"

### A_CcTx_like.png

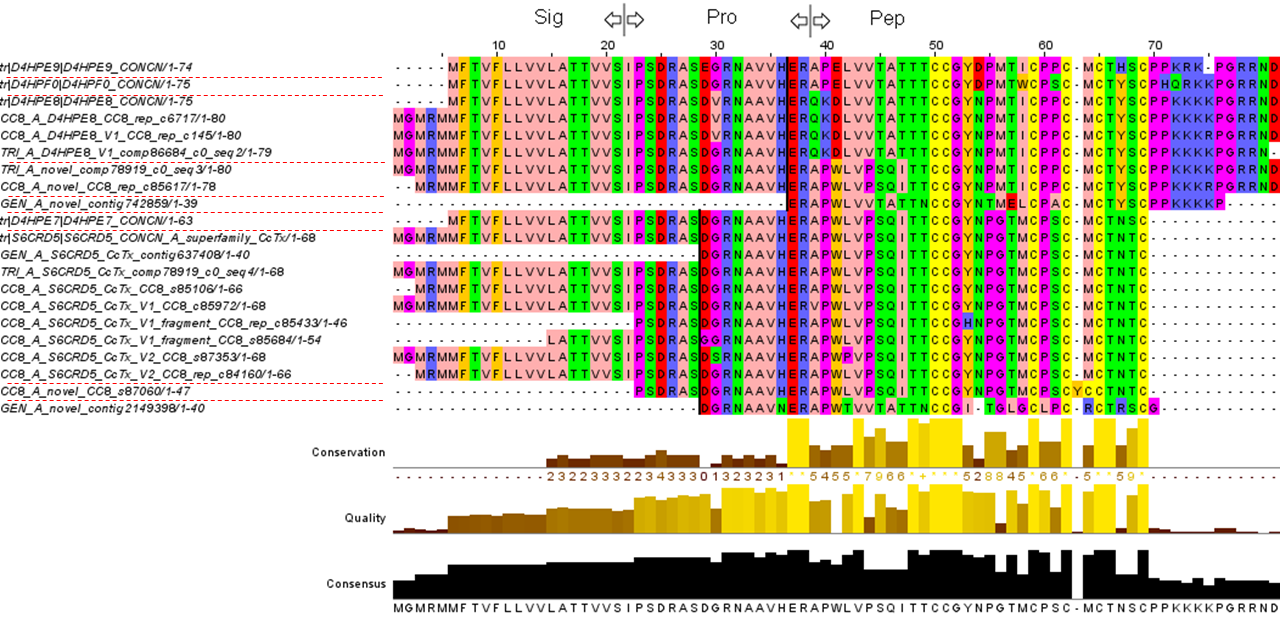

### A_CnI_like.png

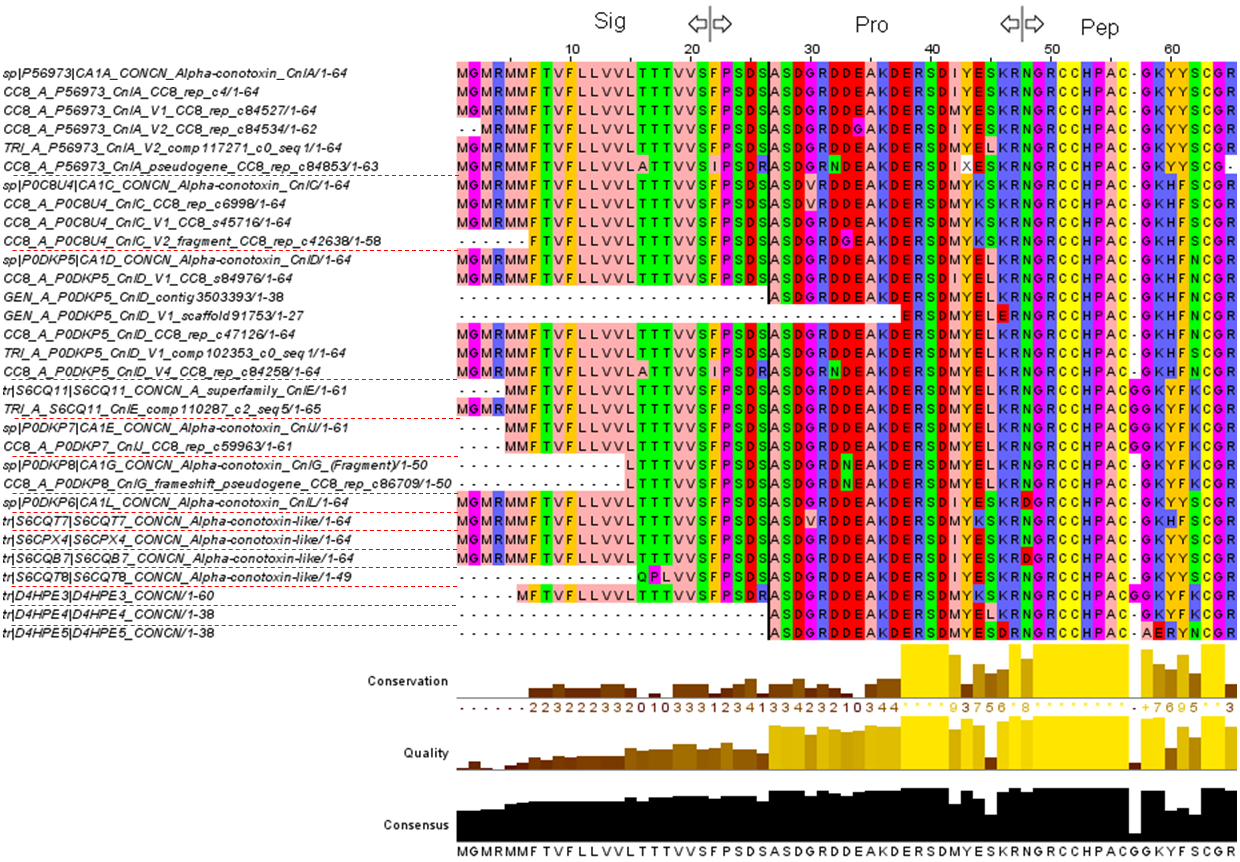

### A_D4HPE6_like.png

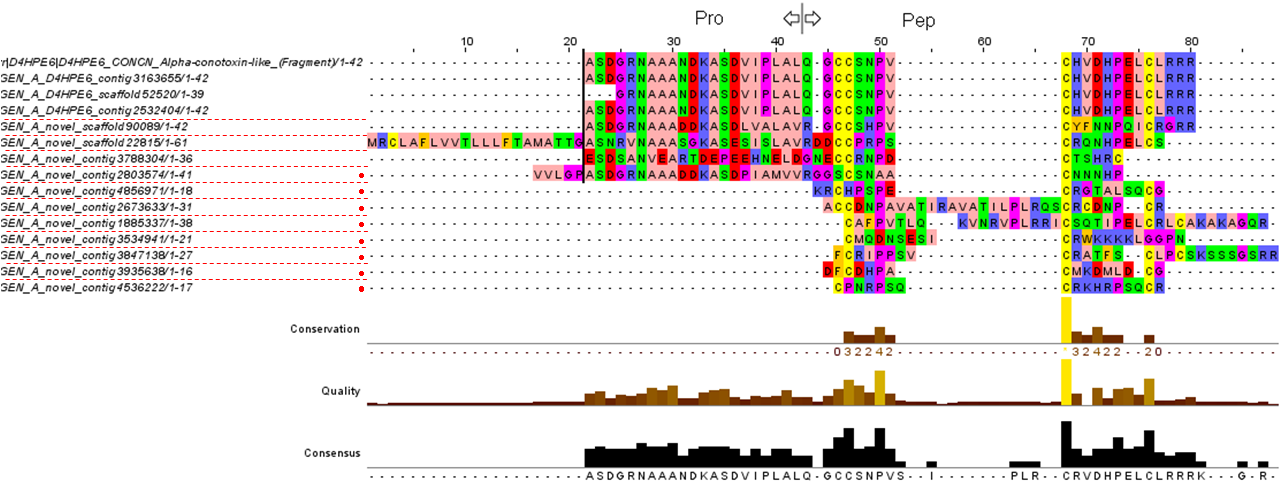

### B_Conantokin.png

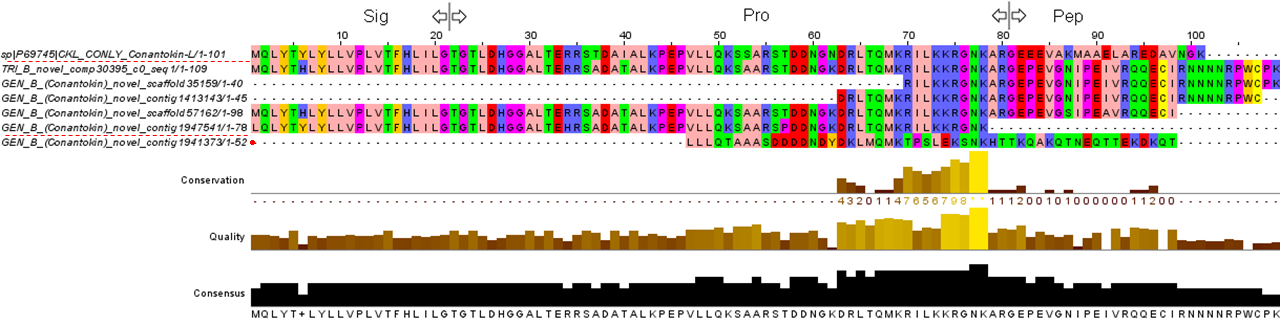

### B_Linear_conopeptide.png

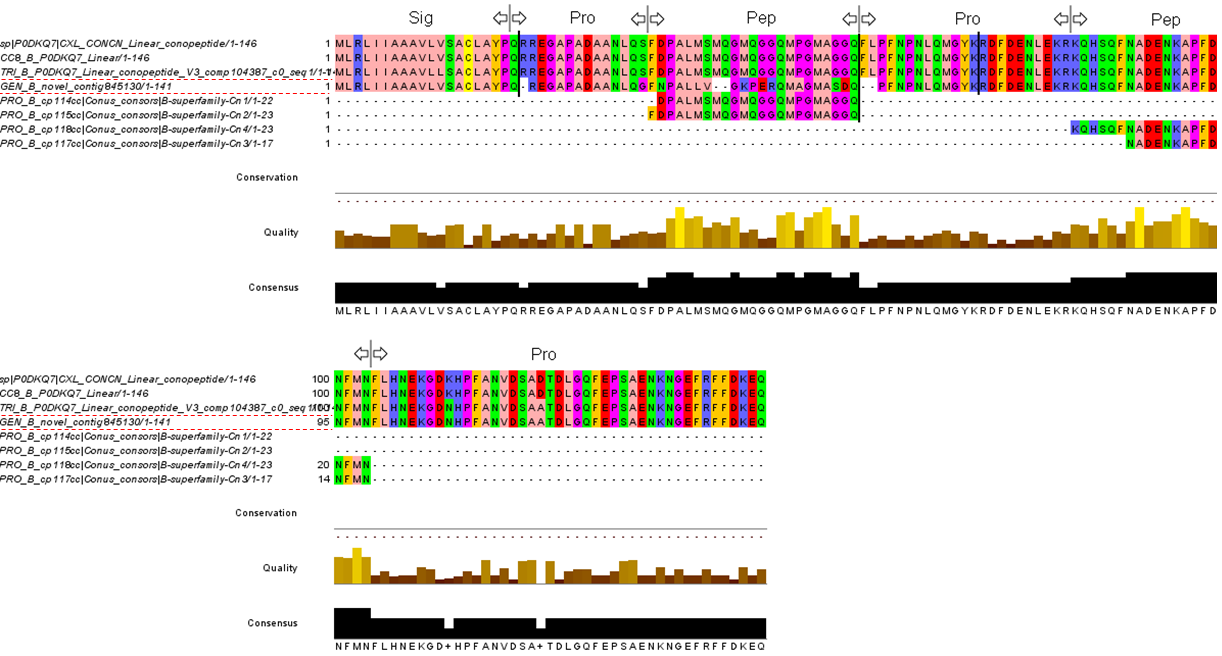

### C_contulakin.png

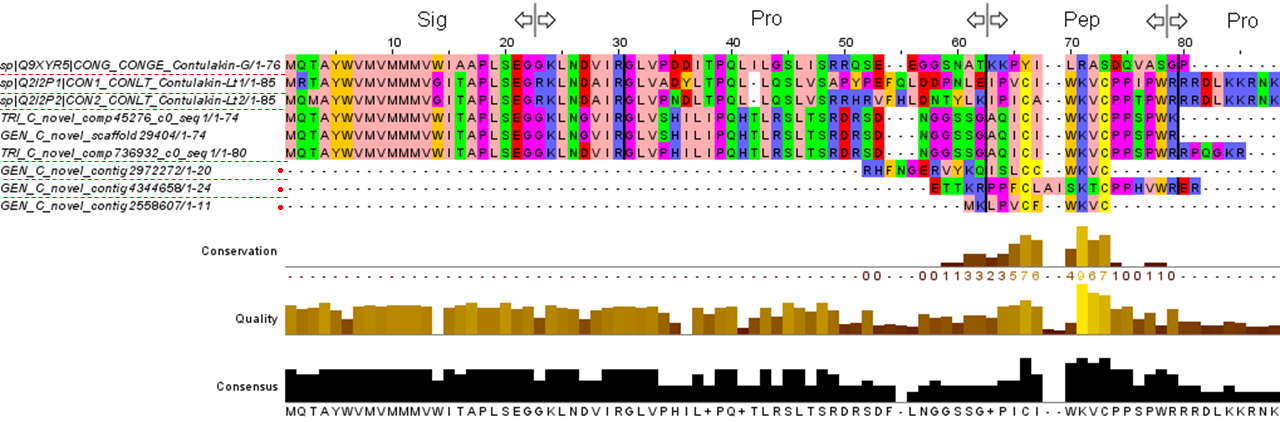

### Conkunitzin.png

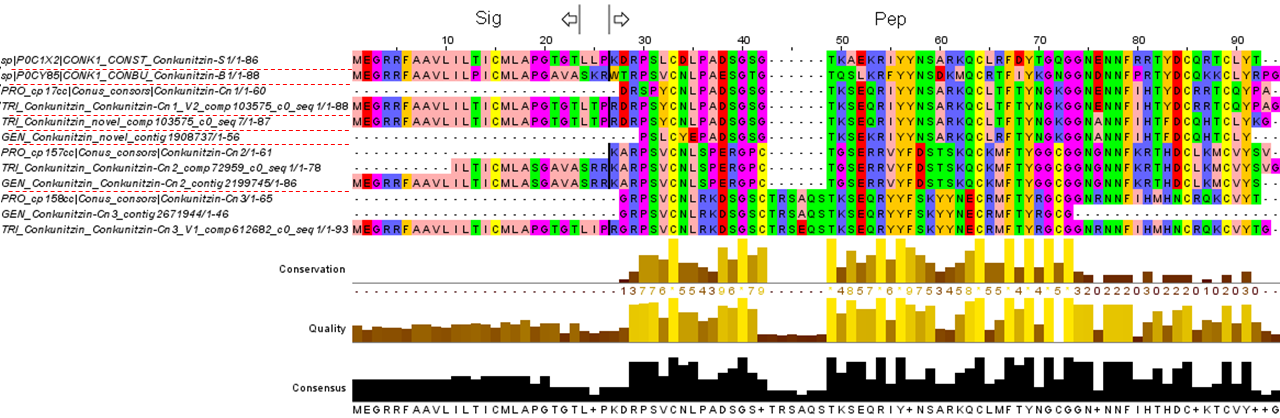

### ConoCAP.png

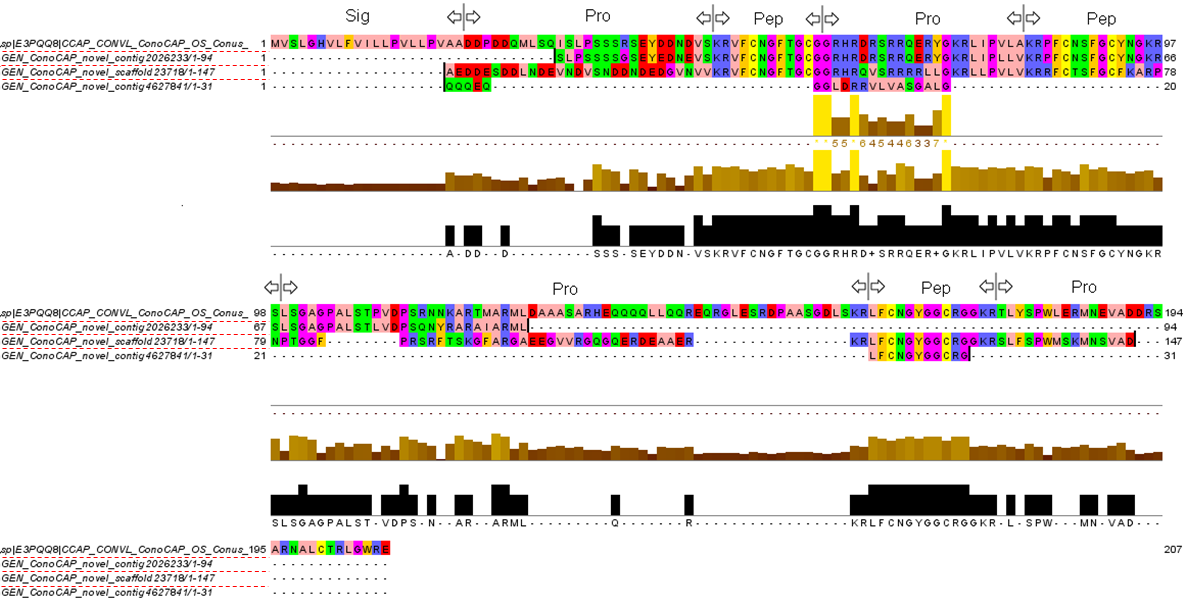

### Conodipine.png

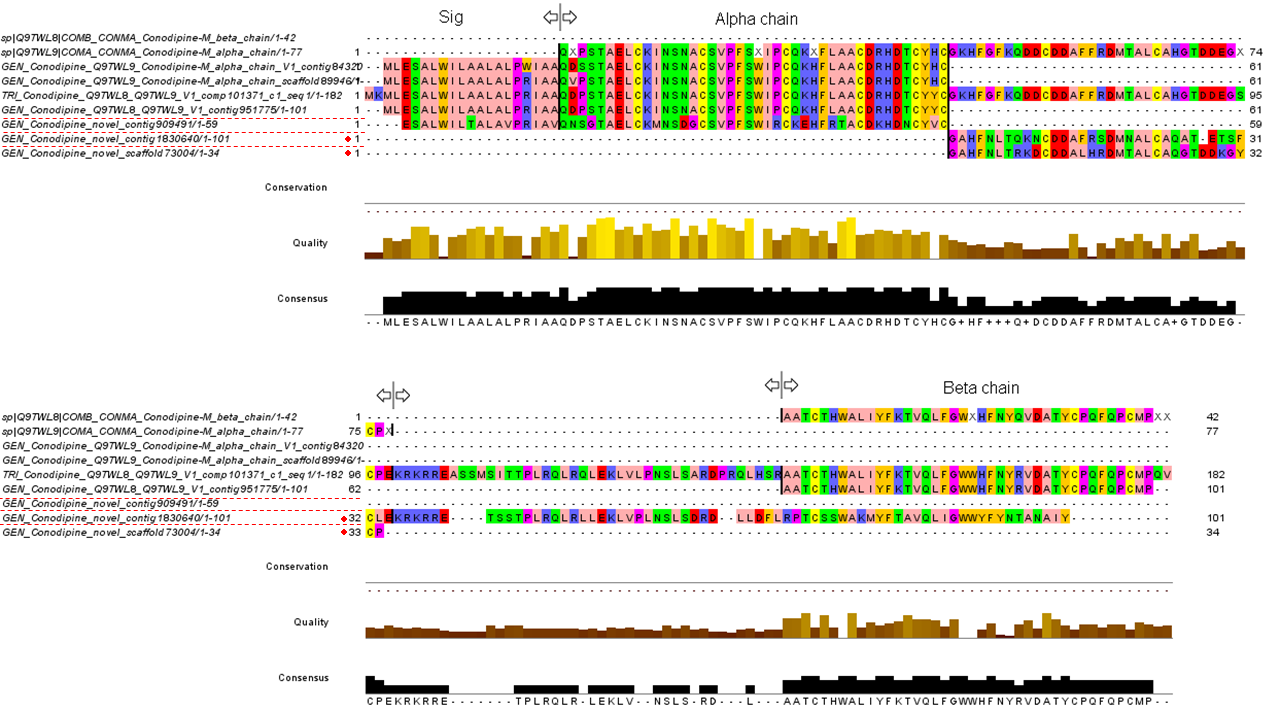

### Conophysin.png

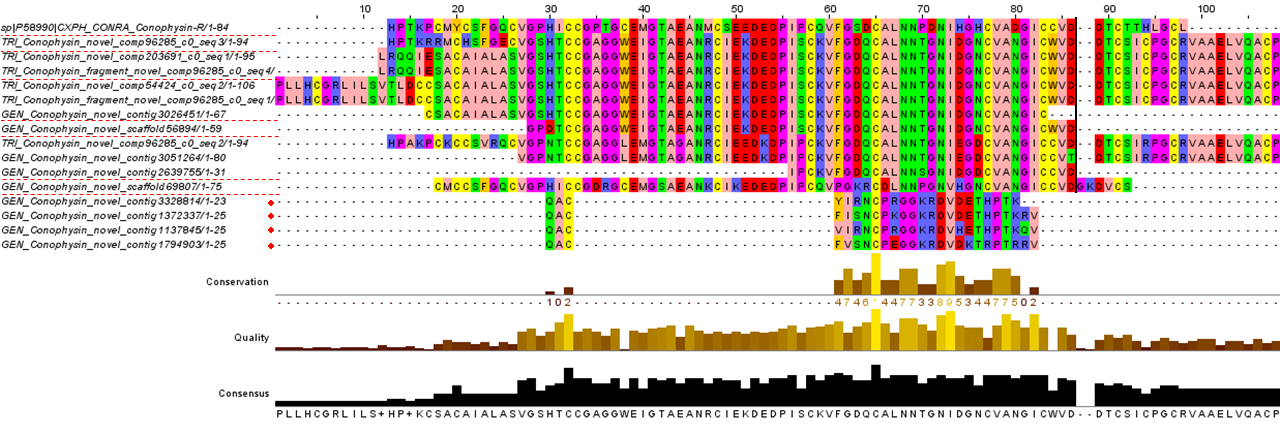

### Conoporin.png

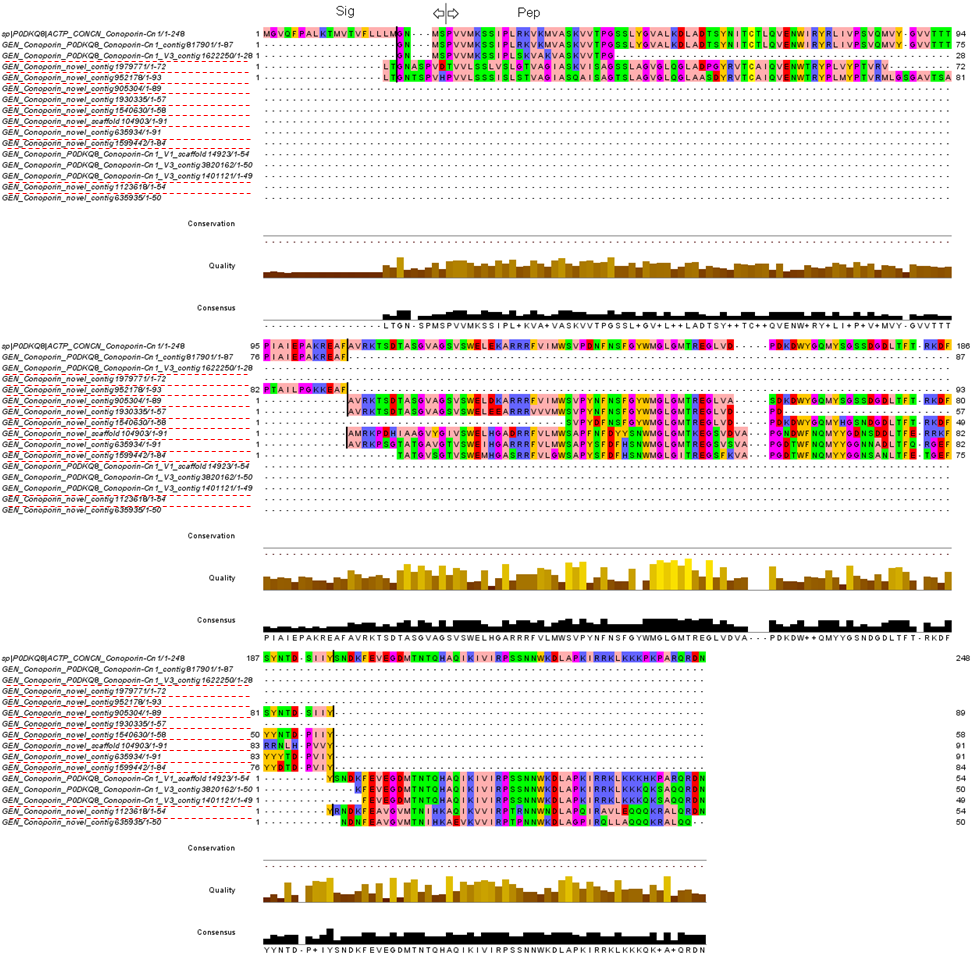

### Contryphan.png

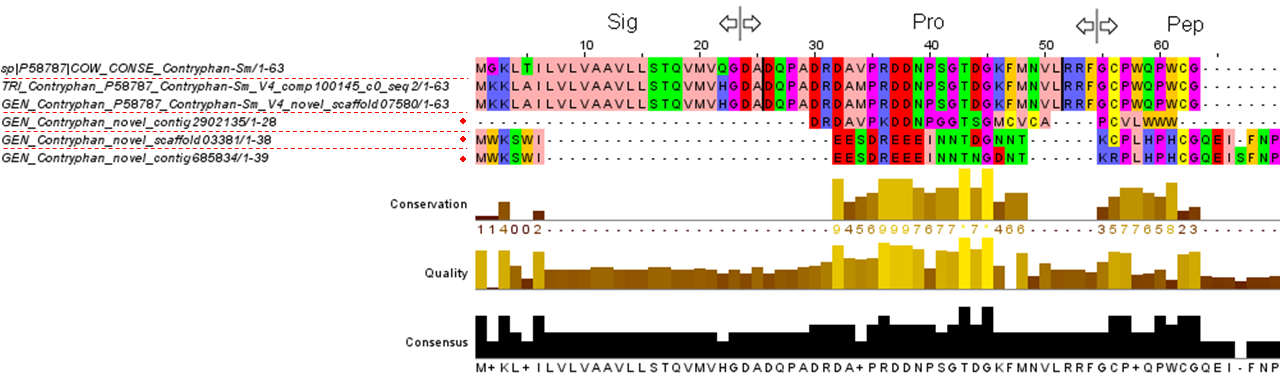

### I1.png

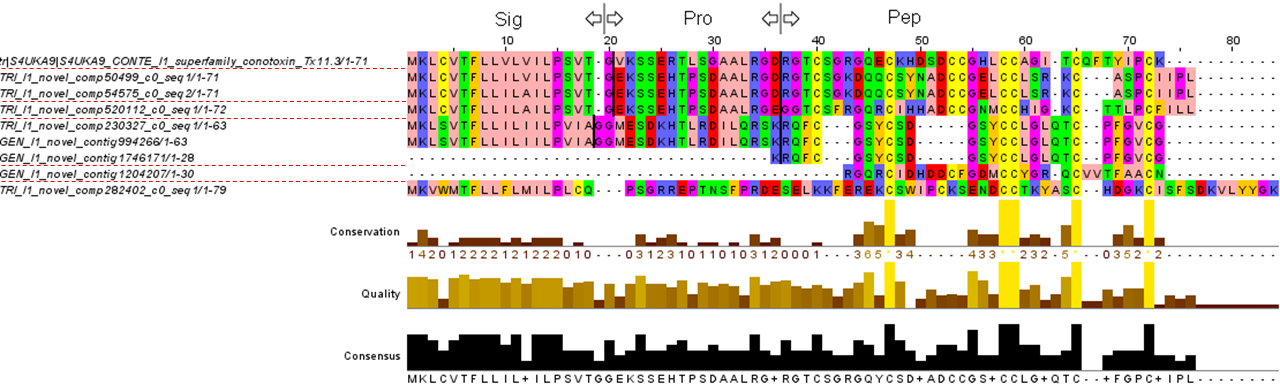

### I2.png

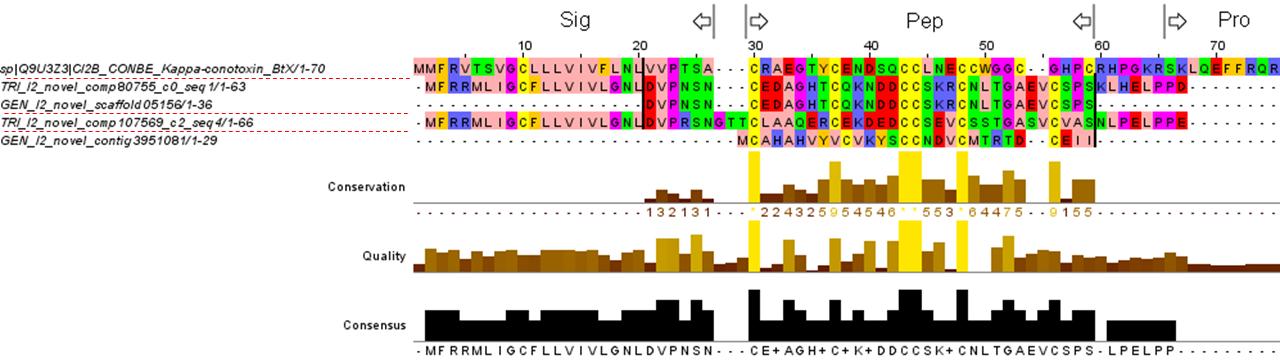

### I3.png

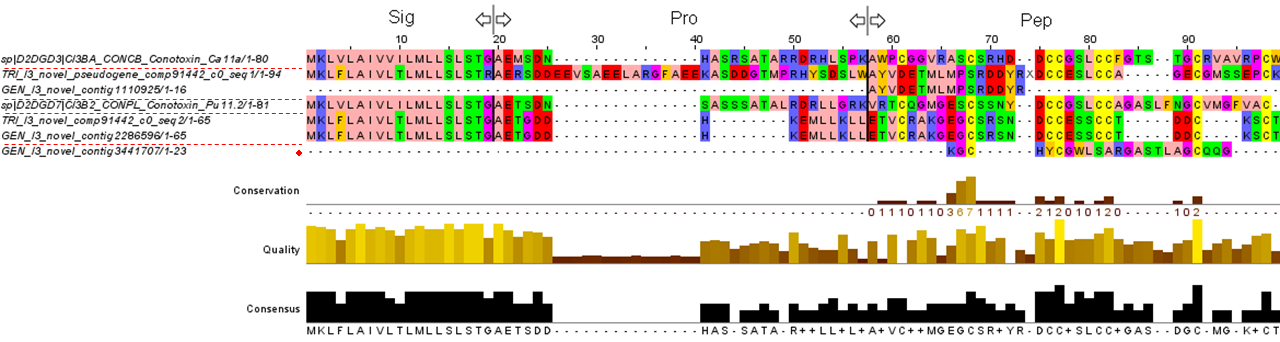

### J.png

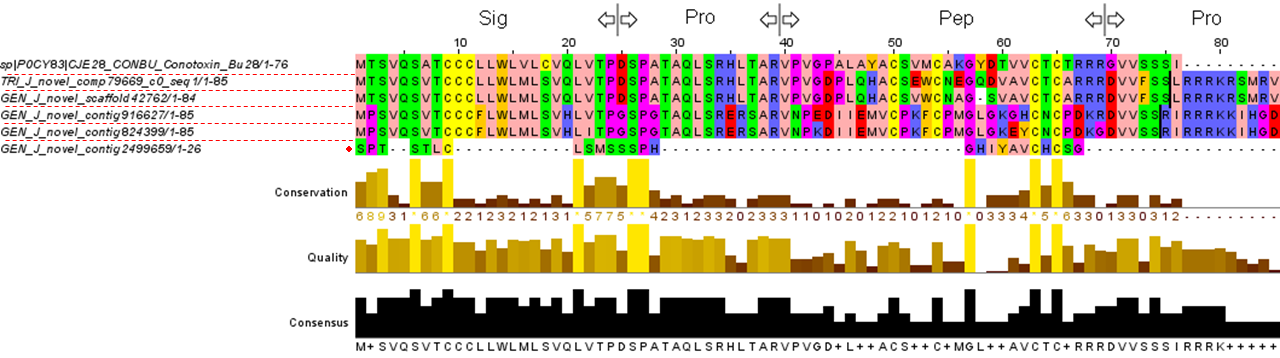

### K.png

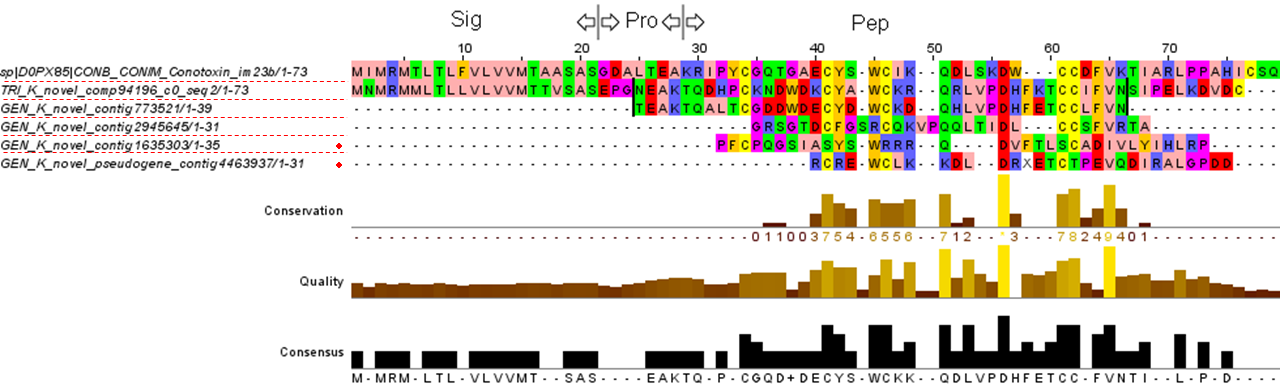

### M_superfamily_CnIII_like.PNG

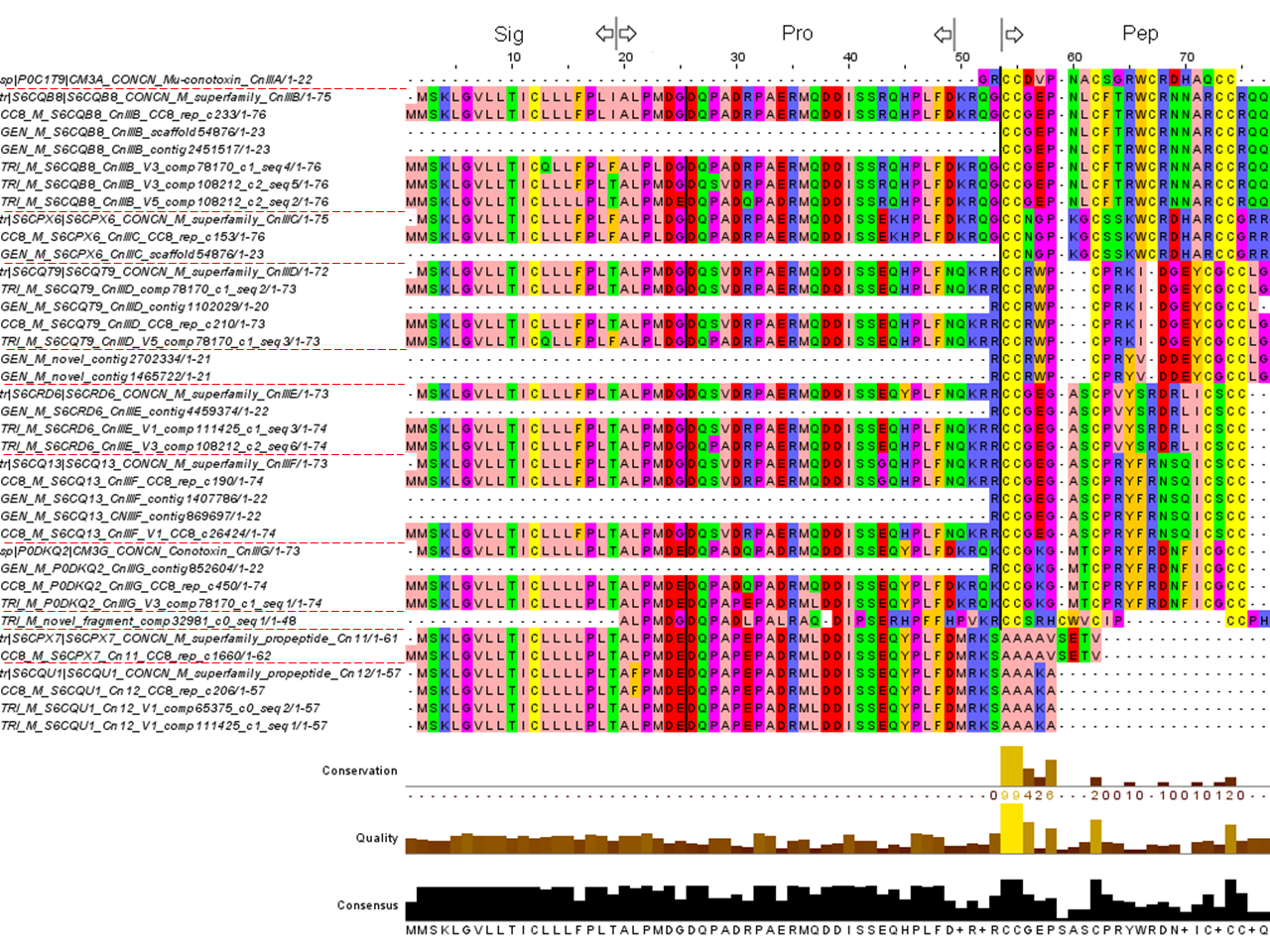

### M_superfamily_Conomarphin_like.PNG

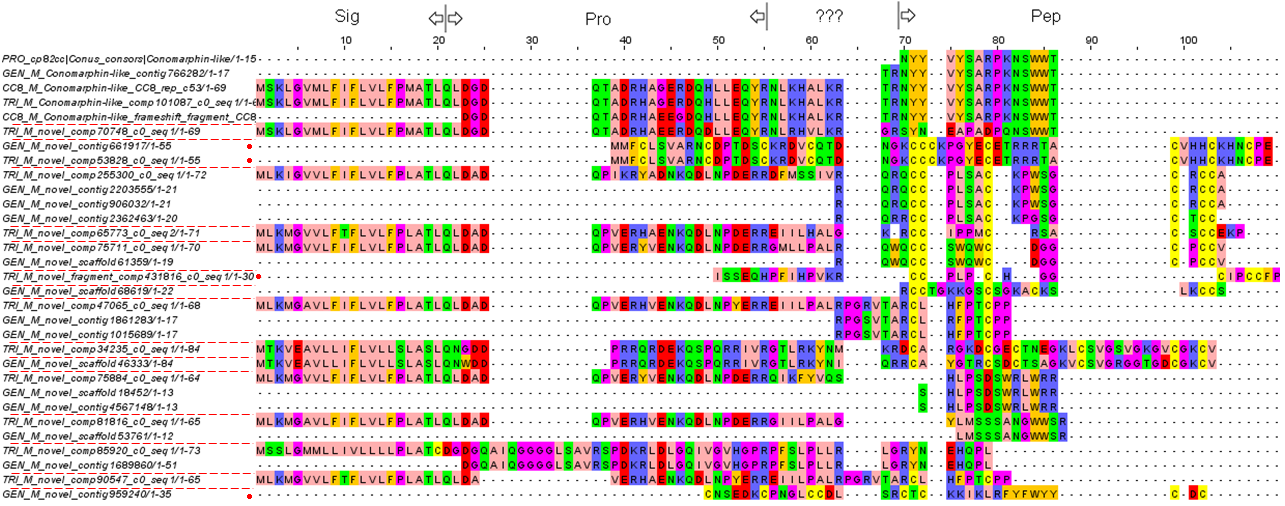

### O1_CnVI_like.png

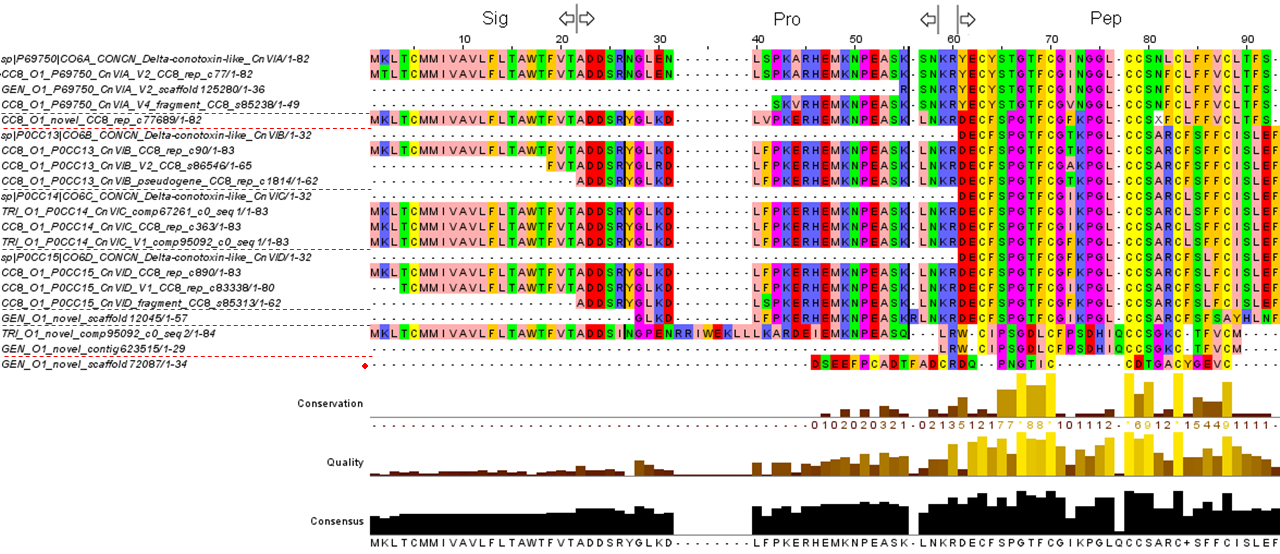

### O1_CnVII_like.png

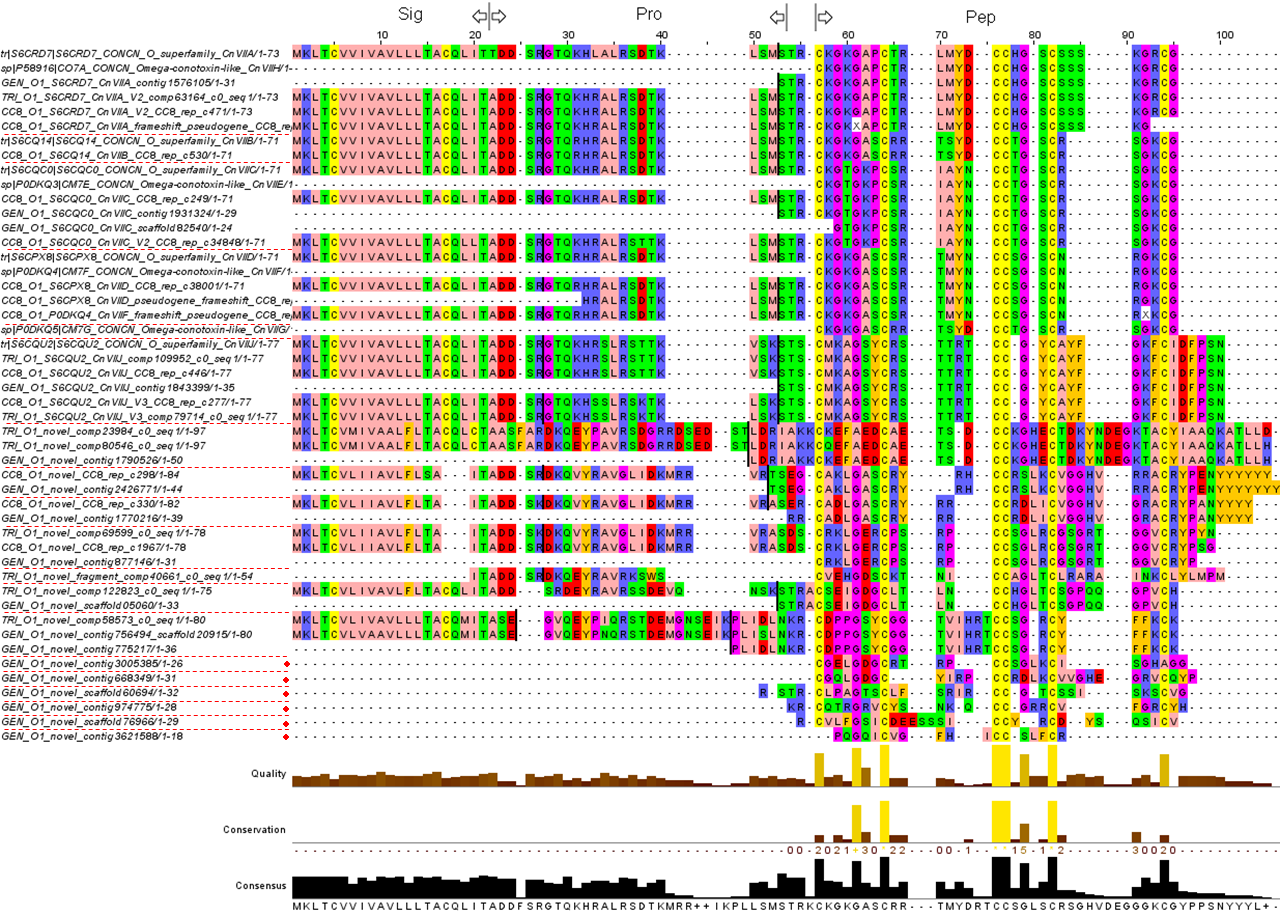

### O2.png

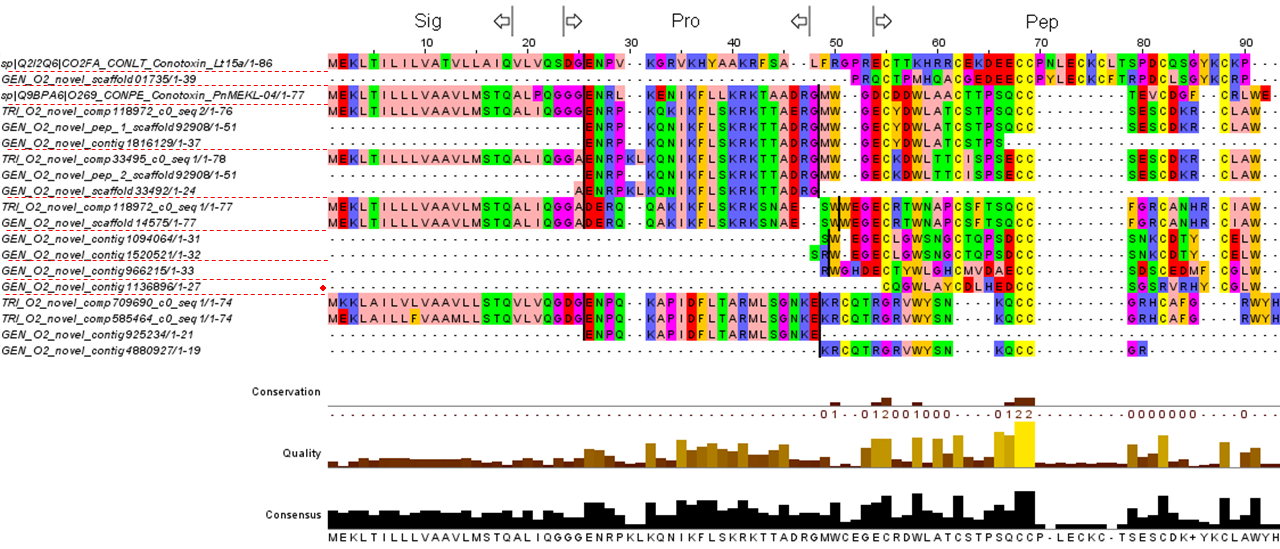

### O3.png

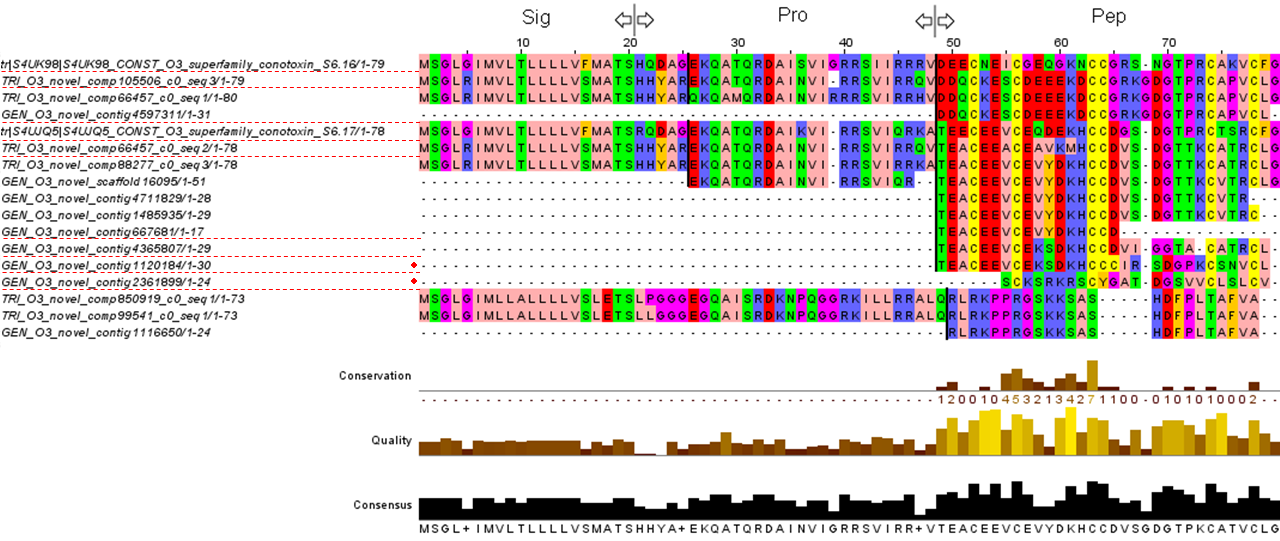

### P.png

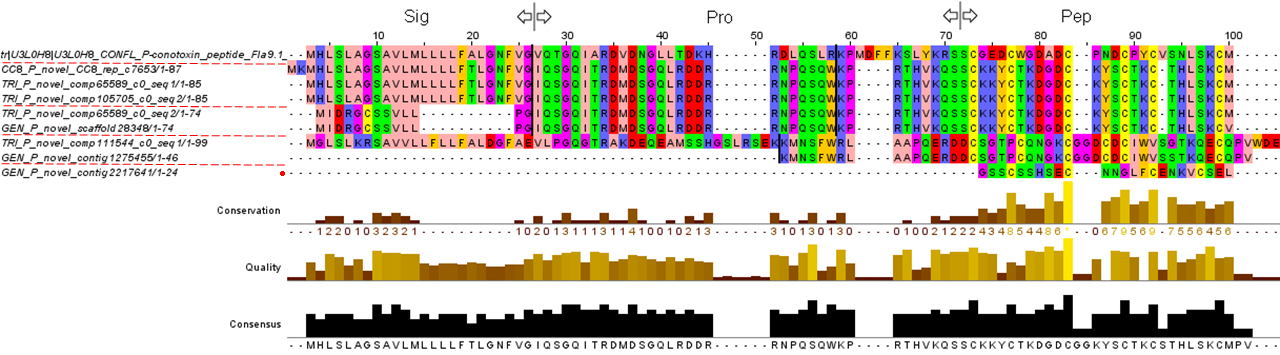

### S_superfamily.PNG

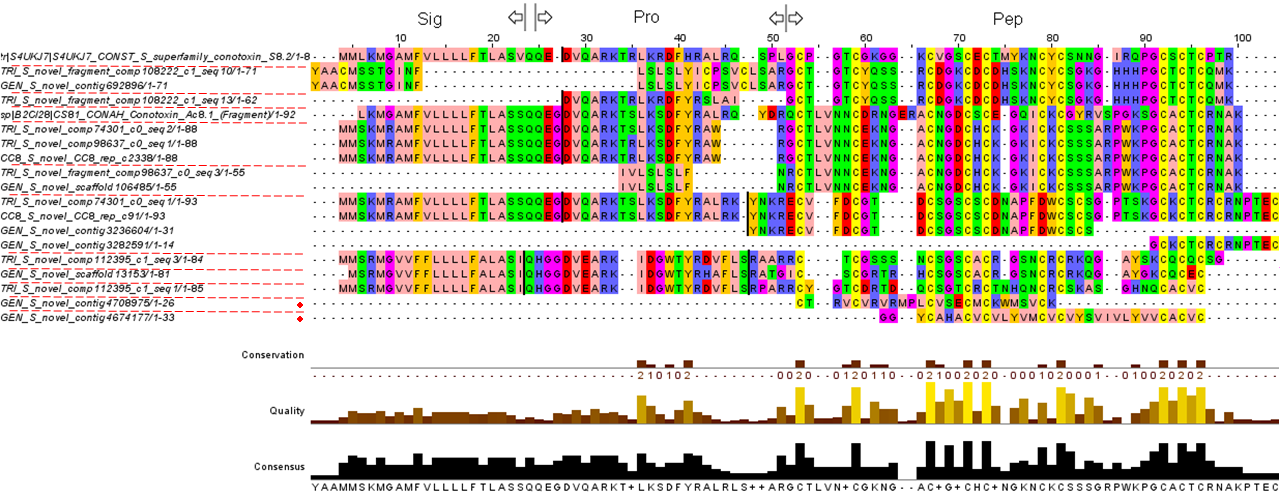

### T.png

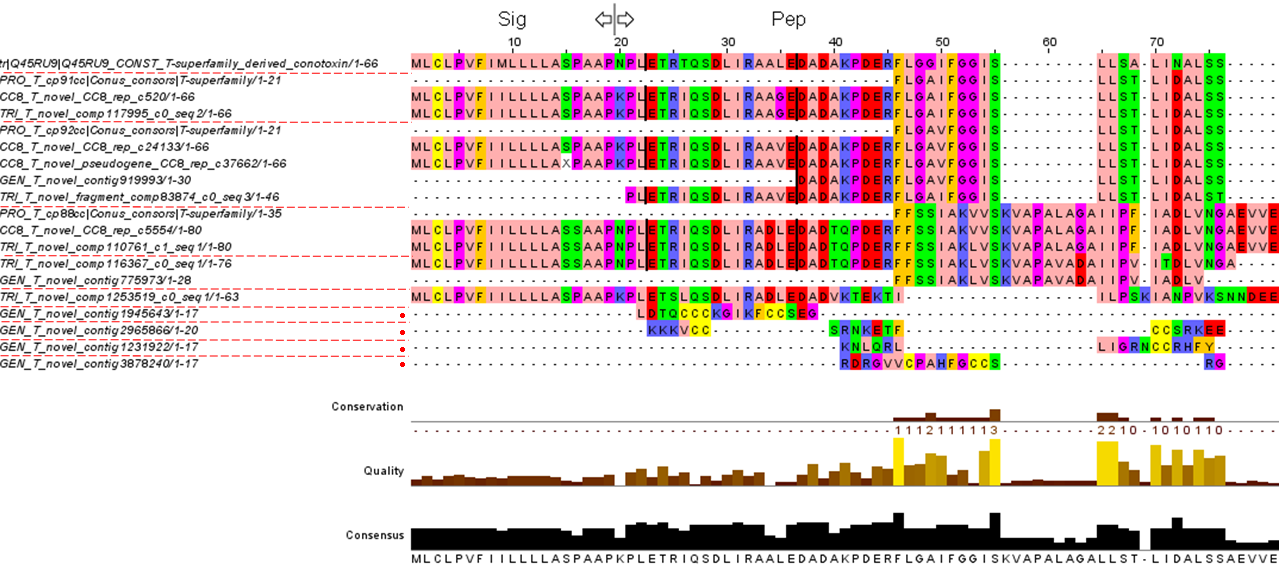

### V.png

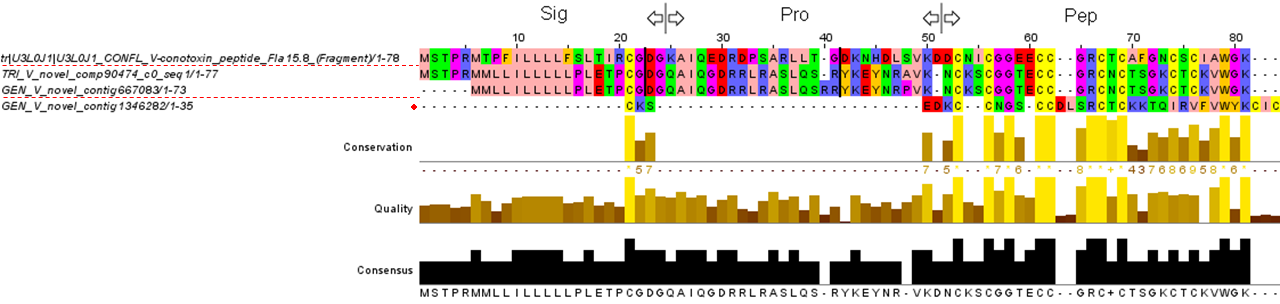
